# Dynamic regulatory module networks for inference of cell type-specific transcriptional networks

**DOI:** 10.1101/2020.07.18.210328

**Authors:** Alireza Fotuhi Siahpirani, Sara Knaack, Deborah Chasman, Morten Seirup, Rupa Sridharan, Ron Stewart, James Thomson, Sushmita Roy

## Abstract

Multi-omic datasets with parallel transcriptomic and epigenomic measurements across time or cell types are becoming increasingly common. However, integrating these data to infer regulatory network dynamics is a major challenge. We present Dynamic Regulatory Module Networks (DRMNs), a novel approach that uses multi-task learning to infer cell type-specific cis-regulatory networks dynamics. Compared to existing approaches, DRMN integrates expression, chromatin state and accessibility, accurately predicts cis-regulators of context-specific expression and models network dynamics across linearly and hierarchically related contexts. We apply DRMN to three dynamic processes of different experimental designs and predict known and novel regulators driving cell type-specific expression patterns.

## Background

Transcriptional regulatory networks connect regulators such as transcription factors to target genes, and specify the context specific patterns of gene expression. Changes in regulatory networks can significantly alter the type or function of a cell, which can affect both normal and disease processes. The regulatory interaction between a transcription factor (TF) and a target gene’s promoter is dependent upon TF binding activity, histone modifications and open chromatin, that have all been associated with cell type-specific expression [1–4]. To probe the dynamic and cell type-specific nature of mammalian regulatory networks, several research groups are generating matched transcriptomic and epigenomic data from short time courses or for cell types related by a branching lineage [5–7]. However, integrating these datasets to infer cell type-specific regulatory networks is an open challenge.

Existing computational methods to infer cell type-specific networks while integrating different types of measurements can be grouped into two main categories: (i) Regression-based methods, (ii) Probabilistic graphical model-based methods. Regression-based methods use linear and non-linear regression to predict mRNA levels as a function of chromatin marks [8, 9] and/or transcription factor occupancies [9] and can infer a predictive model of mRNA for a single condition (time point or cell type). These regression approaches are applied to each context individually and have not been extended to model multiple related time points or cell types, which is important to study networks transition between different time points and cell states. Probabilistic graphical models, namely, dynamic Bayesian networks (DBNs), including input-output Hidden Markov Models [10] and time-varying DBNs [11] have been developed to examine gene expression dynamics with static ChIP-seq datasets. However, both of these approaches are suited for time courses only, and do not accommodate branching structure of cell lineages.

To systematically integrate parallel transcriptomic and epigenomic datasets to predict cell type-specific regulatory networks, we have developed a novel dynamic network reconstruction method, Dynamic Regulatory Module Networks (DRMNs). DRMNs are based on a non-stationary probabilistic graphical model that predicts regulatory networks in a cell type-specific manner by leveraging their relatedness, for example, by time or a lineage. DRMNs represent the cell type regulatory network by a concise set of gene expression modules, defined by groups of genes with similar expression levels, and their associated regulatory programs. The module-based representation of regulatory networks enables us to reduce the number of parameters to be learned and increases the number of samples available for parameter estimation. To learn the regulatory programs of each module at each time point, we use multi-task learning that shares information between related time points or cell types.

We applied DRMNs to four datasets measuring transcriptomic and epigenomic profiles in multiple time points or cell types. Two of these datasets, one microarray [12] and one sequencing, study cellular reprogramming in mouse and measure several histone marks and mRNA levels [7], with the sequencing data additionally measuring accessibility. The third dataset includes RNA-seq and ATAC-seq profiles for mouse dedifferentiation. Finally, the fourth dataset measures transcriptional and epigenomic profiles during early human differentiation to precursor cell types from four main lineages [13]. DRMN learned a modular regulatory program for each of the cell types by integrating chromatin marks, open chromatin, sequence specific motifs and gene expression. Compared to an approach that does not model dependencies among cell types, DRMN was able to predict regulatory networks that reflected the relatedness among the cell types, while maintaining high predictive power of expression. Furthermore, integrating cell type specific chromatin data with cell type invariant sequence motif data enabled us to better predict expression than each data type alone. Comparison of the inferred regulatory networks showed that they change gradually over time, and identified key regulators that are different between the cell types (e.g., Klf4 and Myc in the embryonic stem cell (ESC) state and Six6 and Irx1 in hepatocyte dedifferentiation).Taken together our results show that DRMN is a powerful approach to infer cell type-specific regulatory networks, which enables us to systematically link upstream regulatory programs to gene expression states and to identify regulator and module transitions associated with changes in cell state.

## Results

### Dynamic Regulatory Module Networks (DRMNs)

DRMNs are used to represent and learn context-specific networks, where contexts can be cell types or time points, while leveraging the relationship among the contexts. DRMN’s design is motivated by the difficulty of inferring a context-specific regulatory network from expression when only a small number of samples are available for each context. DRMNs are applicable to different types of contexts; for ease of description we consider cell types as contexts. In a DRMN, the regulatory network for each context is represented compactly by a set of modules, each representing a discrete expression state, and the regulatory program for each module (**Figure 1**). The regulatory program is a predictive model of expression that predicts the expression of genes in a module from upstream regulatory features such as sequence motifs and epigenomic signals. The module-based representations of regulatory networks [14–16], enables DRMN to pool information from multiple genes to learn a predictive regulatory program. In addition, DRMN leverages the relationship between cell types defined by a lineage tree. The modules and regulatory programs are learned simultaneously using a multi-task learning framework to encourage similarity between the models learned for a cell type and its parent in the lineage tree. DRMN allows two ways to share information across cell types: regularized regression (DRMN-FUSED) and graph structure prior (DRMN-ST) (See **Methods**). Both approaches are able to share information across time points effectively, however, the DRMN-FUSED approach is computationally more efficient.

**Figure 1.**
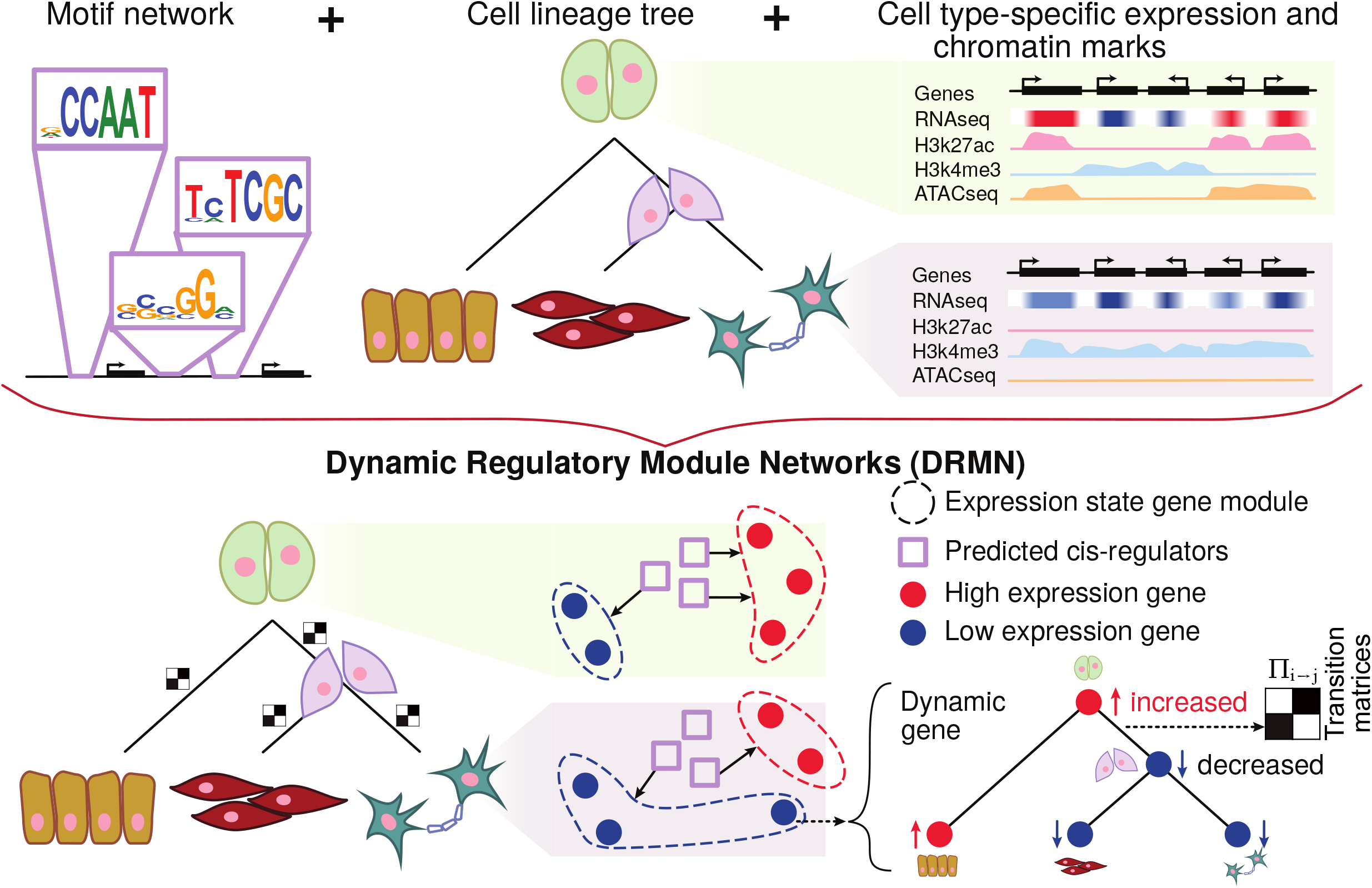
The outline of Dynamic Regulatory Module Networks (DRMN) method. Inputs are a lineage tree over the cell types, cell type-specific expression levels, a shared skeleton regulatory network (*e*.*g*. sequence specific motif network), and optionally cell type-specific features such as histone modification marks or chrmatin accessibility signal. The output is a learned DRMN, which consists of cell type-specific expression state modules, their regulatory programs, and transition matrices describing dynamics between the cell types.

### DRMNs offer a flexible framework to integrate diverse regulatory genomic features

DRMN’s predictive model can be used to incorporate different types of regulatory features, such as sequence-based motif strength, accessibility, and histone marks. We first examined the relative importance of context-specific (e.g., chromatin marks and accessibility) and context independent features (e.g., sequence motifs) for building an accurate gene expression model. We compared the performance of both versions of DRMNs, DRMN-ST and DRMN-FUSED, on different feature sets using an array and a sequencing dataset, both studying mouse cellular reprogramming (**Figure 2**). The array dataset profiles three stages: the starting differentiated cell state (Mouse Embryonic Fibroblast (MEF)), a stalled intermediate stage, (pre-iPSCs) and the end point embryonic cell state (ESC). The sequencing dataset has an additional time point, MEF48 between MEF and pre-iPSCs. Our metric for comparison was Pearson’s correlation between true and predicted expression in each module in a three fold cross-validation setting (**Figure 2A**).

**Figure 2.**
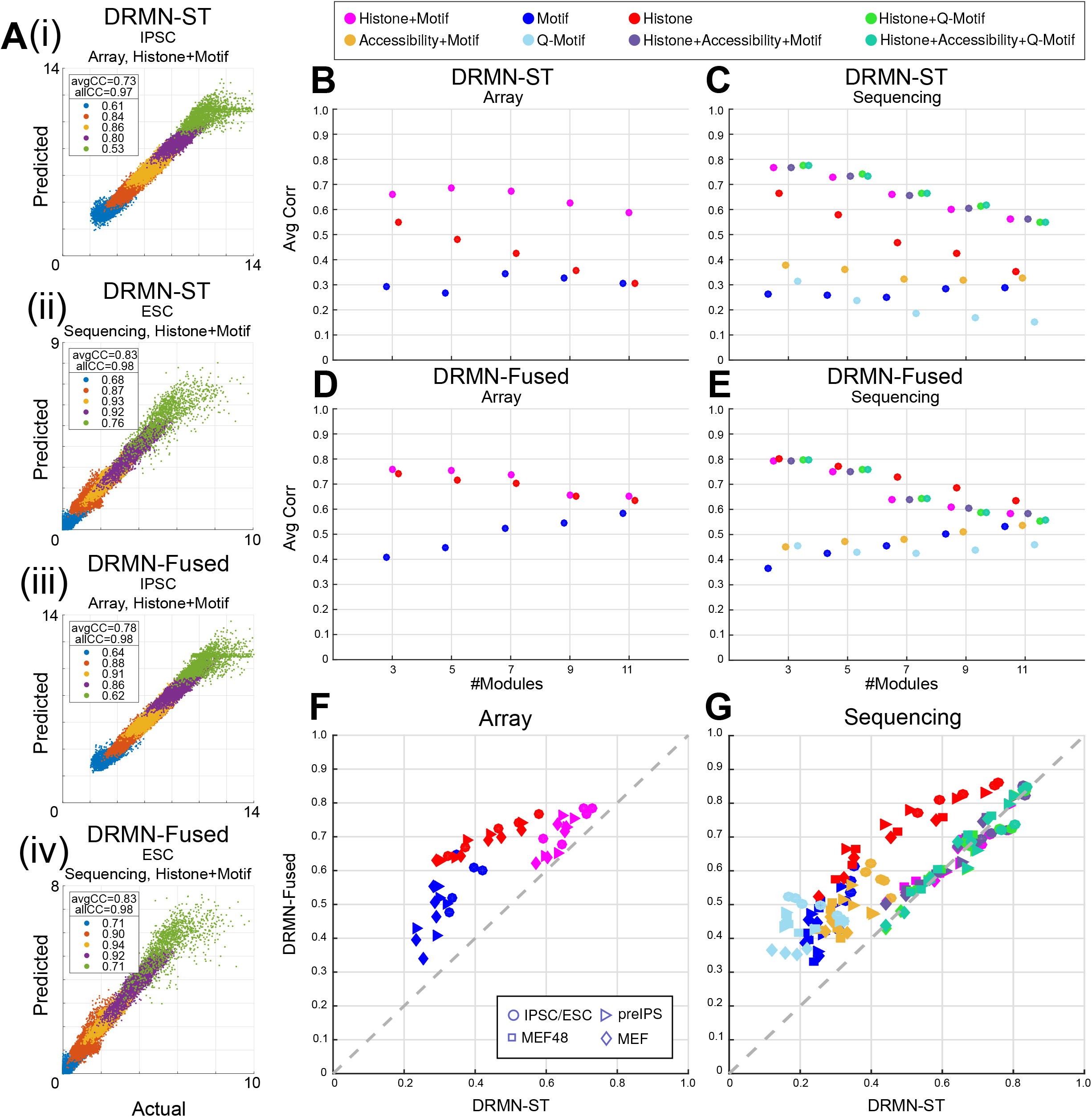
**(A)** Predicted expression *vs*. observed expression, for iPSC/ESC, for **(i)** DRMN-ST on array dataset, **(ii)** DRMN-ST on sequencing dataset, **(iii)** DRMN-FUSED on array dataset, and **(iv)** DRMN-FUSED on sequencing dataset. Average per-module correlation averaged across cell lines as a function of different number of modules, for **(B)** DRMN-ST on array dataset, **(C)** DRMN-ST on sequencing dataset, **(D)** DRMN-FUSED on array dataset, and **(E)** DRMN-FUSED on sequencing dataset. Average per-module correlation for individual cell lines for DRMN-ST *vs*. DRMN-FUSED for **(F)** array dataset, and **(G)** sequencing dataset. Each shape correspond to a cell line and each color correspond to a different feature set.

We first compared DRMN-ST (**Figure 2B,C**) and DRMN-FUSED (**Figure 2D,E**) using sequence-specific motifs alone (Motif), histone marks (Histone) and a combination of the two (Histone+Motif), as these features were available for both array and sequencing datasets. In both models, motifs alone (blue markers) have low predictive power, which is consistent across different *k* and for both array (**Figure 2B, D**) and sequencing (**Figure2C, E**) data. Histone marks alone (red marker) have higher predictive power, however adding both histone marks and motif features (magenta) has the best performance for *k* = 3 and 5, with the improved performance to be more striking for DRMN-ST. For DRMN-Fused, Histone only and Histone+Motif seemed to perform similarly, although at higher *k* using histone marks alone is better. Between different cell types the performance was consistent in array data, while for sequencing data, the MEF and MEF48 cell types were harder to predict than ESC and preIPSC (**Additional file 1 Figure S1**).

We next examined the contribution of accessibility (ATAC-seq) data in predicting expression using the sequencing dataset as accessibility was measured only in this dataset (**Figure 2C, E**). We incorporated the ATAC-seq data in five ways: as a single feature defined by the aggregated accessibility of a particular promoter combined with motifs (Accessibility+motif, orange markers), using ATAC-seq to quantify the strength of a motif instance (Q-Motif, light blue markers), combining the Accessibility feature with histone and motifs (dark purple marker), combining Q-Motif with histone (Histone+Q-motif, light green), and the Accessibility feature with histone and Q-motifs (Histone+Accessibility+Q-motif, dark green, **Figure 2C, E**). We also considered ATAC-seq as a single feature, but this was not very helpful (**Additional file 1 Figure S2**).

Combining the accessibility feature together with motif feature improves performance over the motif feature alone (**Figure 2C, E** orange vs. dark blue markers). The Q-Motif feature (light blue markers) was better than the motif only feature (dark blue) at lower *k* (*k*=3), however, surprisingly, did not outperform the sequence alone features. One possible explanation for this is that the Q-Motif feature is sparser because a motif instance that is not accessible will have a zero value. Finally, we compared the Accessibility feature combined with chromatin and motif features to study additional gain in performance (**Figure 2C, E**, purple markers). Interestingly, even though using accessibility combined with motif features improves the performance over motif alone, addition of ATAC-seq feature to histone marks and motif does not change the performance of either version of DRMN compared to histone and motifs (**Figure 2C**, magenta markers). This suggests that combination of multiple chromatin marks capture the dynamics of expression levels better than the chromatin accessibility signal. It is possible that the overall cell type specific information captured by the accessibility profile is redundant with the large number of chromatin marks in this dataset and we might observe a greater benefit of ATAC-seq if there were fewer or no marks. We observe similar trends with Histone+Q-Motif and Histone+Accesibility+Q-motif features (light and dark green) which perform on par to each other, and close to histone+motif and histone+Accessibility+motif.

DRMN-ST and DRMN-FUSED behaved in a largely consistent way for these different feature combinations with the exception of the Histone+Motif feature where DRMN-Fused was not gaining in performance at higher *k*. When we directly compared DRMN-ST to DRMN-FUSED, DRMN-FUSED was able to outperform DRMN-ST on both Motif and Histone and comparable on Histone+motif on array data (**Figure 2F**). On the sequencing data, DRMN-FUSED had a higher performance that DRMN-ST on Motif, Q-Motif, Accesibilty+motif and Histone alone features (**Figure 2G**). Both models performed similarly when combining Histones with other feature types. It is likely that DRMN-ST learns a sparser model at the cost of predictive power (**Additional file 1 Figure S3**). For the application of DRMNs to real data, we focus on DRMN-FUSED due to its improved performance.

### Multi-task learning approach is beneficial for learning cell type-specific expression patterns

We next assessed the utility of DRMN to share information across cell types or time points while learning predictive models of expression by comparing against several baseline models: (1) those that do not incorporate sharing (RMN), or (2) those that are clustering-based (GMM-Indep and GMM-Merged). GMM-Indep applied Gaussian Mixture Model (GMM) clustering to gene expression values of each cell type independently and GMM-Merged, applied GMM to the merged expression matrix from all cell types. For GMM-Merged, the expression predictions were scored per cell-type.

We evaluated the model performance using quality of predicted expression in a three-fold cross-validation setting, where a model was trained on two-thirds of the genes and used to predict expression for the held-aside one-third of the genes. We ran each experiment over a range of the number of modules, *k ∈* {3, 5, 7, 9, 11}. Expression prediction was evaluated using two metrics. First, the overall expression prediction was assessed using the Pearson’s correlation between actual and predicted expression of genes in the test set. Second, the average Pearson’s correlation of true and predicted expression of genes in the test set in each module. The first metric examines how each model (e.g., expression-based clustering approach versus a predictive model based on regulatory features), explains the overall variation in the data. The second metric assesses the value of using additional regulatory features, such as, sequence and chromatin to predict expression. The clustering-based baselines offer simple approaches to describe the major expression patterns across time but link *cis*-regulatory elements to expression changes only as post-processing steps. For both DRMN and RMN, we learned predictive models of expression in each module using different regulatory feature sets (**Figure 3, Additional file 1 Figure S4**): motif alone (Motif), histone marks alone (Histone) and combing motif and histone marks (Histone+Motif). We performed these experiments on the array and sequencing datasets for mouse reprogramming.

**Figure 3.**
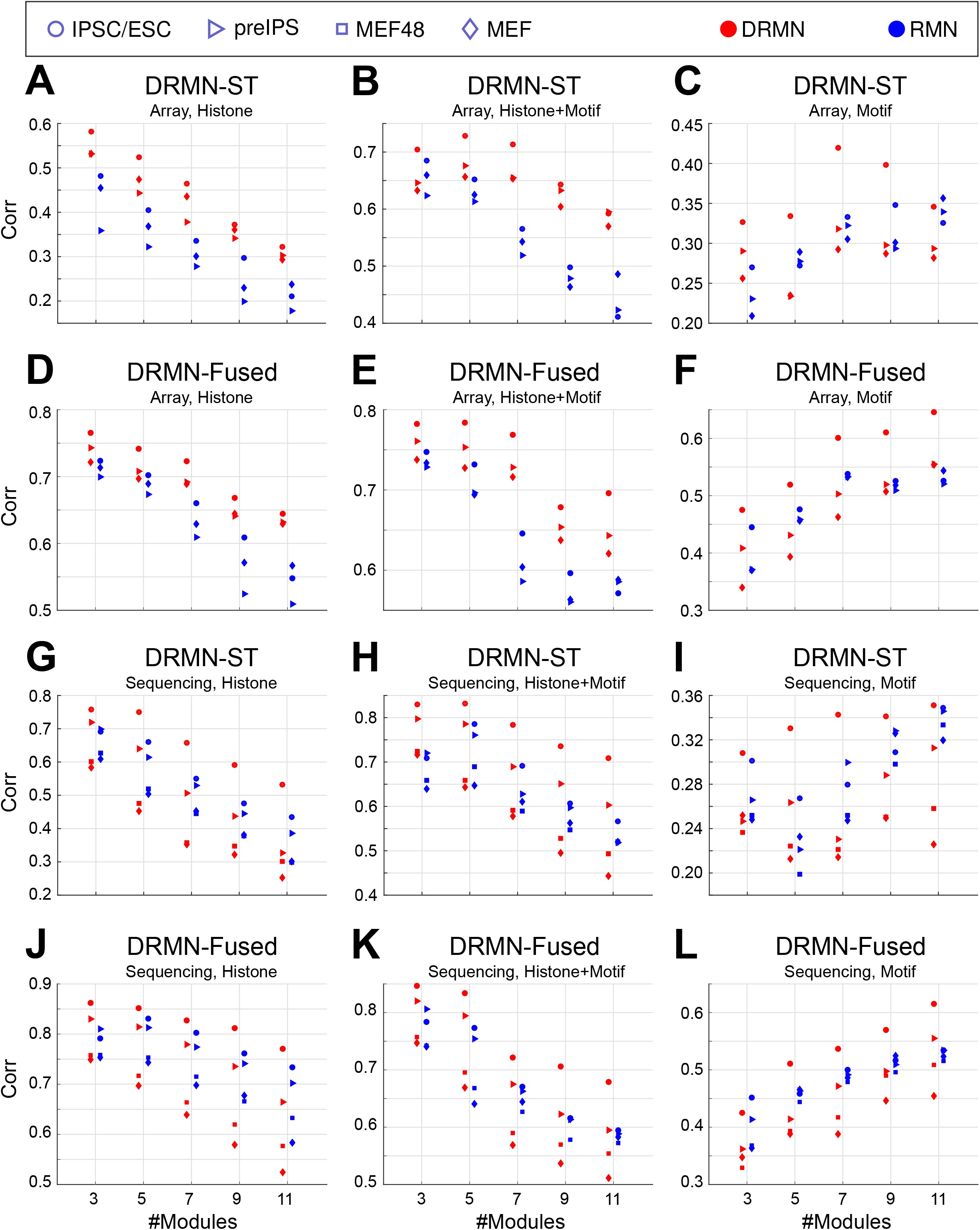
Average per-module correlation for individual cell lines as a function of different number of modules for single task and multi task versions of the method, for **A-C)** DRMN-ST on sequeincing dataset, **D-F)** DRMN-FUSED on sequencing dataset, **G-I)** DRMN-ST on array dataset, and **J-L)** DRMN-FUSED on array dataset. Each shape correspond to a cell line and each color correspond to a different method.

When comparing DRMNs to RMNs on array data, both versions of DRMN outperform corresponding RMN versions on histone only and histone + motif features, (**Figure 3A-F**), blue for DRMN and red for RMN). On motifs, the difference between the models was dependent upon the cell line and the number of modules, *k*. In particular, both DRMN-Fused and DRMN-ST outperformed RMN on the iPSC/ESC state, and was similar for pre-iPSC and MEF when comparing across different *k*. On sequencing data (**Figure 3G-L**), both DRMN-Fused and DRMN-ST were better than RMNs for most *k* in ESC and pre-IPSCs when considering Histone and Histone+Motifs. When using motifs, the performance depended upon the DRMN implementation and *k*. In particular, DRMN-Fused was at par or better than RMNs for the other cell lines. DRMN-ST was at par or better than RMNs for most *k*, however we observed decrease in performance in the MEF and MEF48 cell lines for higher *k* (*k* = 9, 11).

We next compared DRMNs and RMNs to the clustering based methods using the two metrics of overall correlation across all genes and per module correlation. When using overall correlation, both variants of DRMNs, RMNs and GMM-Indep, vastly outperform GMM-Merged (**Additional file 1 Figure S4A-F**). This suggests that the gene partitions are likely different between the different cell types and imposing a single structure for all three, as done in GMM-Merged, misses out on the cell type specific aspects of the data. DRMN models performed at par with RMN and GMM-Indep for most cases (**Additional file 1 Figure S4A-F**), with the exception of Histone and Histone+Motif for sequencing data (**Additional file 1 Figure S4A, B**) where GMM-Indep is worse for lower *k*s. These results suggest that based on overall correlation, learning predictive or clustering models for each cell line or cell type is more advantageous than learning a single clustering model (e.g., GMM-Merged), which can better capture the cell type-specific aspects of the data.

We next compared the different models on the basis of the per-module expression levels (**Additional file 1 Figure S4G-L**). DRMN and RMN clearly outperform the GMM-based approaches, which is because GMM clustering produces one value per module and does not capture the within-module variation. The overall high correlation for DRMN and RMN demonstrates that a predictive modeling approach has advantages over a clustering approach as the learned models from the *cis*-regulatory features provide a more fine-tuned model of expression variation. Furthermore, predicting expression across multiple cell types simultaneously using multi-task learning helps to further improve the predictive power of these models.

### Using DRMN to gain insight into regulatory programs of cellular reprogramming

We applied DRMN to gain insights into the regulatory programs of cellular reprogramming from mouse embryonic fibroblasts (MEFs) to induced pluripotent stem cells (iPSCs). We focus on the results obtained on the sequencing dataset for reprogramming from Chronis et al [7] (**Figure 4**), which measured chromatin marks (via ChIP-seq), accessibility (via ATAC-seq) and gene expression (via RNA-seq). Many of the trends are captured in the array data too (**Additional file 1 Figure S5**). DRMN modules learned on the sequencing data exhibit seven distinct patterns of expression in each of the four cellular stages (**Figure 4A**). While the expression patterns remains the same, the number of genes in each module in each cell type varies (**Figure 4A**). We compared the extent of similarity of matched modules between cell types and found that the modules were on average 30-90% similar, exhibiting the lowest similarity at the MEF48 to pre-iPSC transition and the highest between MEF and MEF48 (**Figure 4B**). The repressed modules 1,2 and 3 were less conserved across all cell types compared to the more highly expressed modules. For the array data as well, we observed the greatest dissimilarity between the MEF and pre-iPSC transition (**Additional file 1 Figure S5**). This agrees with the pre-iPSC state exhibiting a major change in transcriptional status during reprogramming.

**Figure 4.**
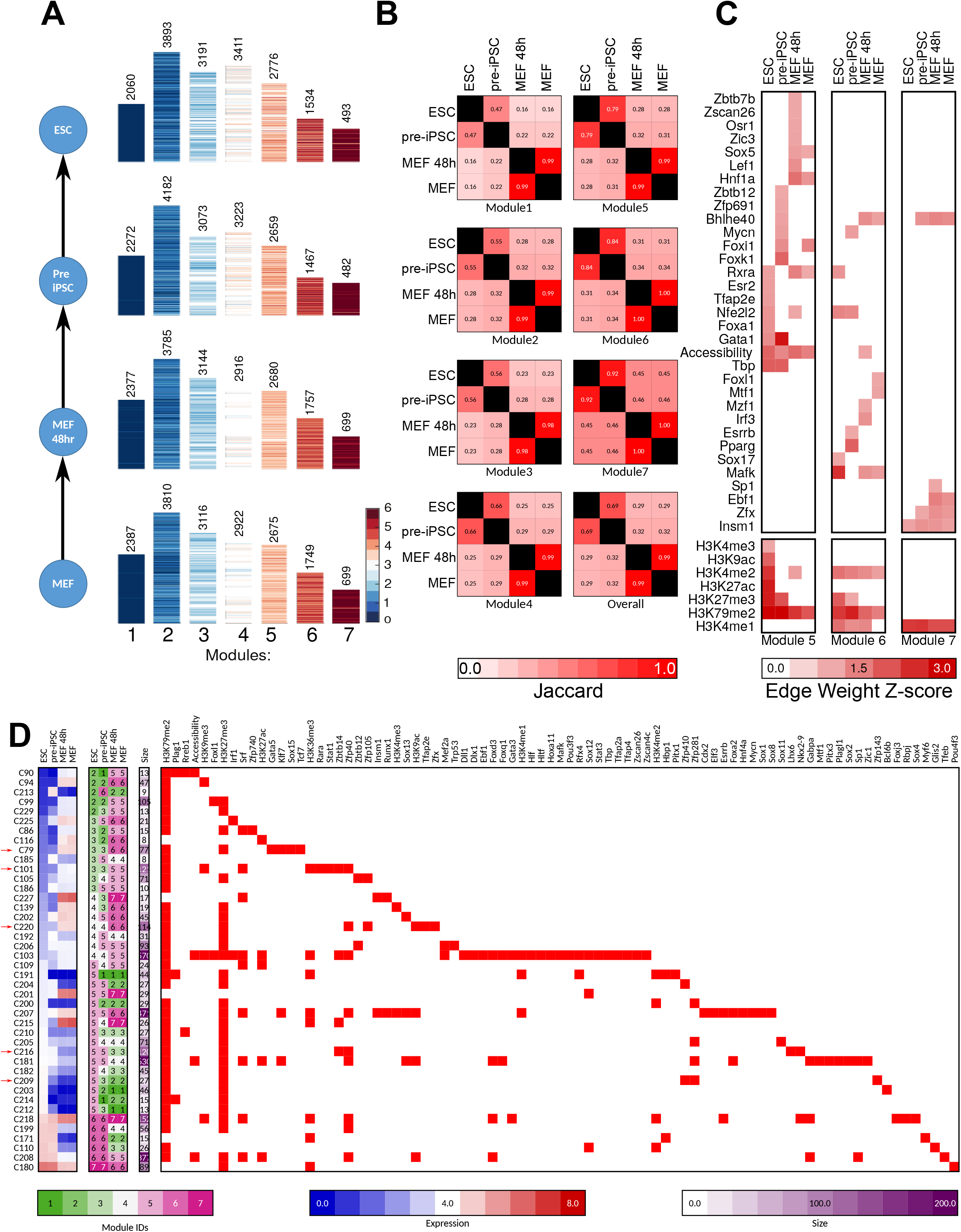
Application of DRMNs to the cellular reprogramming sequencing dataset using histone marks, accessibility and Q-motifs for the feature set. **A**. Show are gene expression patterns of the *k* = 7 modules. The number above the heatmap is the number of genes in that module. **B**. Similarity of modules across cellular stages as measured by F-score. The color intensity is proportional to the match. **C**. Inferred regulatory program for the most highly expressed modules across cell stages. **D**. Transitioning gene sets exhibiting changes into the high expression modules (3, 4, 5). Shown are the mean expression levels of genes in the gene set (left, red-blue heat map), the module assignment (second), the number of genes in each module (third) and the set of regulators for each gene set (white-red heat map). Red arrows depict gene sets discussed in the text.

To interpret the modules, we first tested them for enrichment of Gene Ontology (GO) processes using a FDR corrected hyper-geometric test (P-value *<*0.001). Consistent with the low conservation of genes present in the modules 1-3, we found few process terms that shared enrichment across these modules. In contrast modules 4, 5, and 6 exhibited greater shared enrichment and included house-keeping function such as ribosome biogenesis and general metabolic processes. Several processes were unique to each cell stage. For example in the repressed modules 1 and 2, we saw enrichment of inflammation response to be down-regulated in iPSC, pre-iPSC, while we observed down-regulation of sexual reproduction and gametogenesis in the MEF/MEF48 stages (**Additional file 2**). Processes that were specifically up-regulated in the iPSC, preiPSC stages included the cell-cycle, while muscle development and cell adhesion processes were associated with MEF, MEF48. Overall, the presence of up-regulated cell cycle processes in the ESC/pre-iPSC and muscle related processes in MEF/MEF48 is consistent with the stages.

We next examined the regulatory programs inferred for each module (**Figure 4C, Methods**), and found several known and novel regulatory features associated with each module. For ease of interpretation, we focused on the three modules associated with highest expression (5, 6 and 7). Several histone marks (H3K79me2 and H3K4me1) and accessibility were selected as predictive features for all four cellular stages. In contrast, the transcription factors (TFs) were more specific to each state with a few exceptions (e.g., Insm1 and Bhlhe40). Importantly, we found that several TFs have known roles in the stage in which they are found significant, e.g., Nfe2fl2 in ESC and pre-IPSC module 6, which is known to play an important role in the embryonic stem cell state [17], and Mycn and Esrrb in the pre-IPSC module 6, which are both known to be important for early embryonic stages. We also found several muscle and mesodermal factors associated with MEF (Foxl1 [18], Mtf1 [19] in module 6), MEF48 (Osr1 [20], Left1 [21] in module 5), and both MEF and MEF48 (Sox5 [22] in module 5).

To gain insight into the regulatory features most important for driving transcriptional dynamics, we identified genes that change their module assignments (module transition) between cell stages. Of the 17,358 genes, 11,152 genes changed their module assignments. We clustered them into sets of 5 or more genes and defined 111 gene sets consisting of 10,194 genes (the remaining were singleton or genes with similarity to less than 4 genes). These transitioning gene sets provide insight into the different classes of expression dynamics exhibited by genes. We next predicted regulators, including both chromatin marks and TFs, for these gene sets using a regularized regression model (see **Methods**) and were able to associate regulators with high confidence to 85 gene sets. These gene sets exhibited a variety of transitions, from induced to repressed expression levels and vice-versa from MEF to ESC cell state. We next examined individual gene sets focusing on 42 gene sets with transitions into the high expression modules, 5, 6 or 7 (**Figure 4D**, for transitioning gene sets in array dataset see **Additional file 1 Figure S6**, the full set of gene sets is in **Additional file 3**). In particular, two gene sets (C101 and C220, **Figure 5A, B**) exhibited low expression in ESCs and pre-iPSC and high expression in MEF and MEF48, while another showed the opposite pattern (C216, **Figure 5F**). For the majority of these genesets, we found a combination of TFs and histone marks predicted to regulate them. Among these were factors such Srf which was shown to facilitate reprogramming [23], and Plag1, which is involved in cancer and growth processes and was shown to effect embryonic development [24]. Taken together, application of DRMN to this dataset characterized the dynamic patterns of expression and predicted regulators that could be important for cellular reprogramming and cell state maintenance.

**Figure 5.**
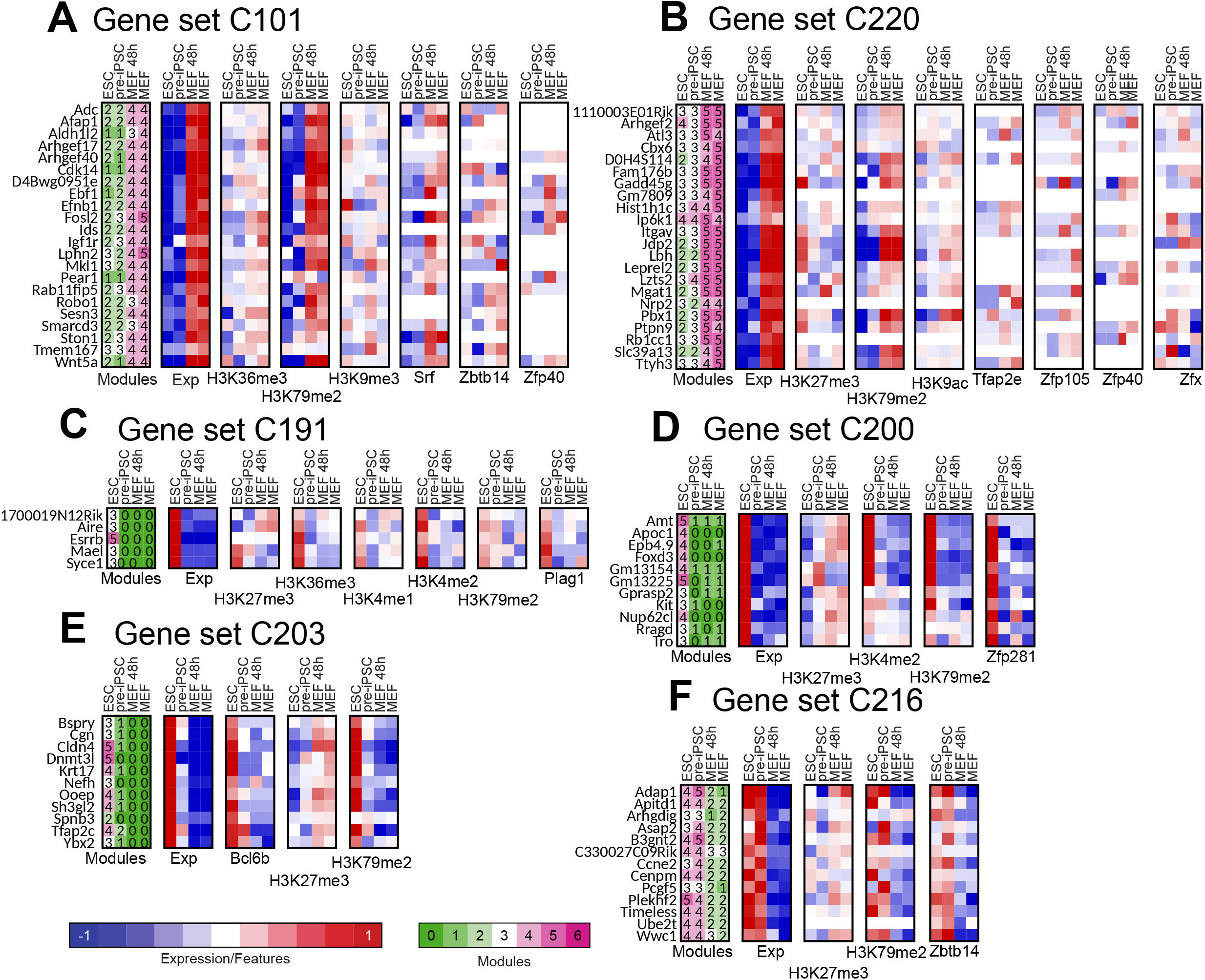
Selected transitioning gene sets in the cellular reprogramming dataset. Each panel shows the member genes of a transitioning gene set (label on top). The columns show the module assignment of each gene, followed by its expression level in each cell type (Exp). The subsequent groups of columns are the levels of the regulator on the gene promoters. The name of the regulator is specified at the bottom of the heat map. All significant regulators are shown. **A-B**. Gene sets exhibiting down-regulation in ESC/pre-iPSC and up-regulation in MEF/MEF48 (Geneset C101, C220). **C-E**. Up-regulation in ESC/iPSC alone (Gene sets C191, C200, C203). **F**. Upregulation in ESC/iPSC and pre-iPSC and down regulation in MEF and MEF48 (Geneset C216).

### Using DRMNs to gain insight into regulatory program dynamics across a long time course

While the reprogramming study demonstrated the application of DRMNs to a short time course (three-four time points), we next tested the ability of DRMN to analyze temporal dynamics of a longer time course with dozens of time points. In particular, we applied DRMNs to examine temporal dynamics of regulatory programs during hepatocyte dedifferentiation. A major challenge in studying primary cells in culture such as liver hepatocytes is that they dedifferentiate from their differentiated state. Maintaining hepatocytes in their differentiated state is important for studying normal liver function as well as for liver-related diseases [25]. Dedifferentiation could be due to the changes in the regulatory program over time, however, little is known about the transcriptional and epigenetic changes during this process. To measure transcriptional dynamics during dedifferentiation, Seirup et al. generated an RNA-seq and ATAC-seq timecourse dataset of 16 time points from 0 hours to 36 hours [26]. We applied DRMN with *k* = 5 modules to this dataset and partitioned genes into five levels of expression at all time points with 1 representing the lowest level of expression and 5 the highest level of expression (**Figure 6A**). Comparison of module assignments across time showed that module 1, associated with the lowest expression was most conserved across time points, while modules 2, 3 and 4, exhibited transitions happening between 4 and 6 hrs and 14 and 16 hrs (**Figure 6B**). The relatively high conservation of the repressed module was in striking contrast to what we observed for the most repressed module in the reprogramming study, where it was least conserved.

**Figure 6.**
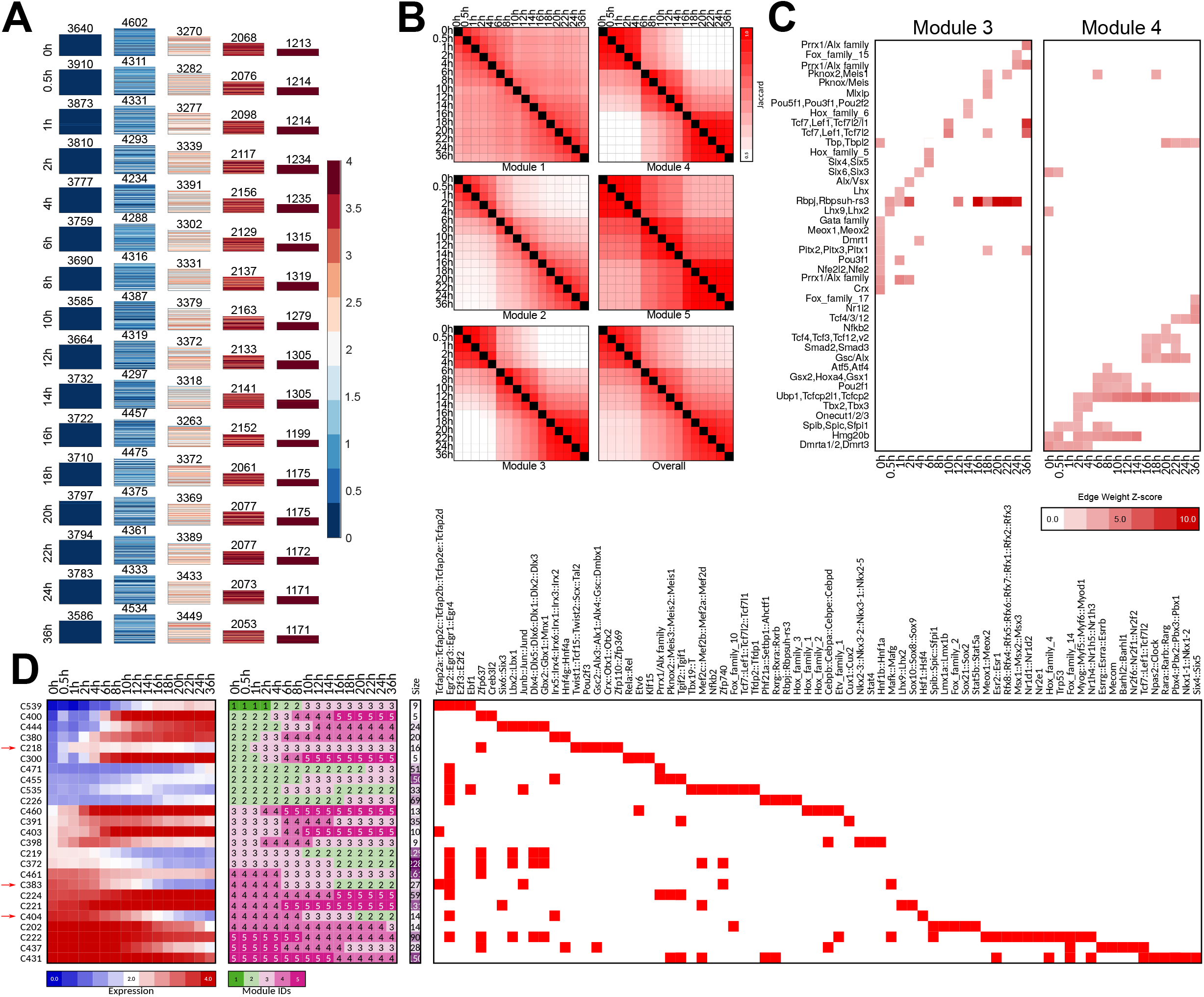
Application of DRMNs to the hepatocyte dedifferentiation dataset using accessibility and Q-motifs for the feature set. **A**. Show are gene expression patterns of *k* = 5 modules for each time point (major row). The number above the heatmap is the number of genes in that module. **B**. Similarity of modules across time points as measured by F-score. The color intensity is proportional to the match. **C**. Inferred regulators for each modules across time. Only modules with highest expression and that had TFs as regulators (3 and 4) are shown. **D**. Transitioning gene sets exhibiting changes into the high expression modules (3, 4, 5). Shown are the mean expression levels of genes in the gene set (left, red-blue heat map), the module assignment (second), the number of genes in each module (third) and the set of regulators for each gene set (white-red heat map). Red arrows depict gene sets discussed in the text.

As before, we tested the genes in each module for GO process enrichment (**Additional file 2**) and found that the repressed module (Module 1) was enriched for developmental processes while the other modules were enriched for diverse metabolic processes. Of these modules 4 and 5, which are associated with higher expression are enriched for more liver-specific metabolic function such as co-enzyme metabolism, acetyl CoA metabolism and modules 2 and 3 were enriched for general housekeeping function such as DNA and nucleic acid metabolism. We next examined the regulators associated with each module. Focusing on the three highly expressed modules (modules 3, 4 and 5), we found several liver factors, e.g., Nfkb [27], Foxk1 and Foxk2 [28, 29], Onecut2 [30, 31], Lhx2 [32] that are likely important for maintenance of the hepatocyte state (**Figure 6C**). In addition, we found several regulators involved in cell fate decision making e.g., Tcf3 [33] and Smad2 [34]. The cell fate regulators are enriched in the later part of the time course indicating their potential roles in dedifferentiation.

To gain insight into fine-grained transition dynamics, we next used the DRMN module assignments to identify genes that transition from one module to another as a function of time, similar to the reprogramming study. Of the 14,793 genes, there were 5,762 genes that change their module assignments. We identified a total of 150 transitioning gene sets spanning 5,762 genes (**Additional file 3**). Many of the transitions were between modules that are adjacent to each other based on expression levels, suggesting that the majority of the dynamic transitions are subtle (e.g., module 4 and 5, **Figure 6D**). However, there were several gene sets that exhibited more drastic transitions, e.g., C404 and C383 transitioning from induced to repressed expression while C218 exhibiting the opposite pattern.

We next predicted regulators for these transitioning gene sets using a regularized Group Lasso model, Multi-Task Group LASSO (MTG-LASSO, **Methods**). This approach was different from what we had for reprogramming and modeled individual gene’s profiles while incorporating their membership in a gene set. Briefly, this approach solves a multi-task learning problem where each task predicts the expression of one gene in the transitioning gene set using regression. Instead of learning a vector of regression weights for each task independently, this approach learns a matrix of coefficients which is regularized such that the same set of regulators are selected for each gene but with different values. Using this approach we identified 84 gene sets with predicted regulators at *≥*60% confidence (**Methods, Additional file 4**). We focused on those gene sets with a transition into modules 3, 4 and 5 and identified a total of 25 gene sets with varying types of transitions (**Figure 6D**). Several of these gene sets exhibited a gradual upregulation of expression, e.g., C444, C400, C218, C380 (**Figure 7**), including transitions at 4 and 6 hrs which had the largest change in module assignment. Regulators associated with these gene sets included pluripotency or developmental regulators, such as Irx1 [35] (C380, C455), Meis3 (C224, C455) [36], Alx1 [37] (C455, C224), and the Egr family (C224). Early growth response (Egr) factors have been shown to play important roles in different liver-related functions including repair and injury [38]. We also identified gene sets with down-regulation of expression, e.g., C437. Key regulators predicted for these gene sets included Nuclear receptor 2 family (Nr2f2, Nr2f6 and Nr2f1), and Hepatocyte nuclear factors (Hnf4g, Hnf4a) that have been implicated in liver-specific function [39]. Taken together, these results suggest that DRMNs can systematically integrate gene expression and accessibility measurements across long time courses and predict both known and novel regulators associated with the dynamics of the process.

**Figure 7.**
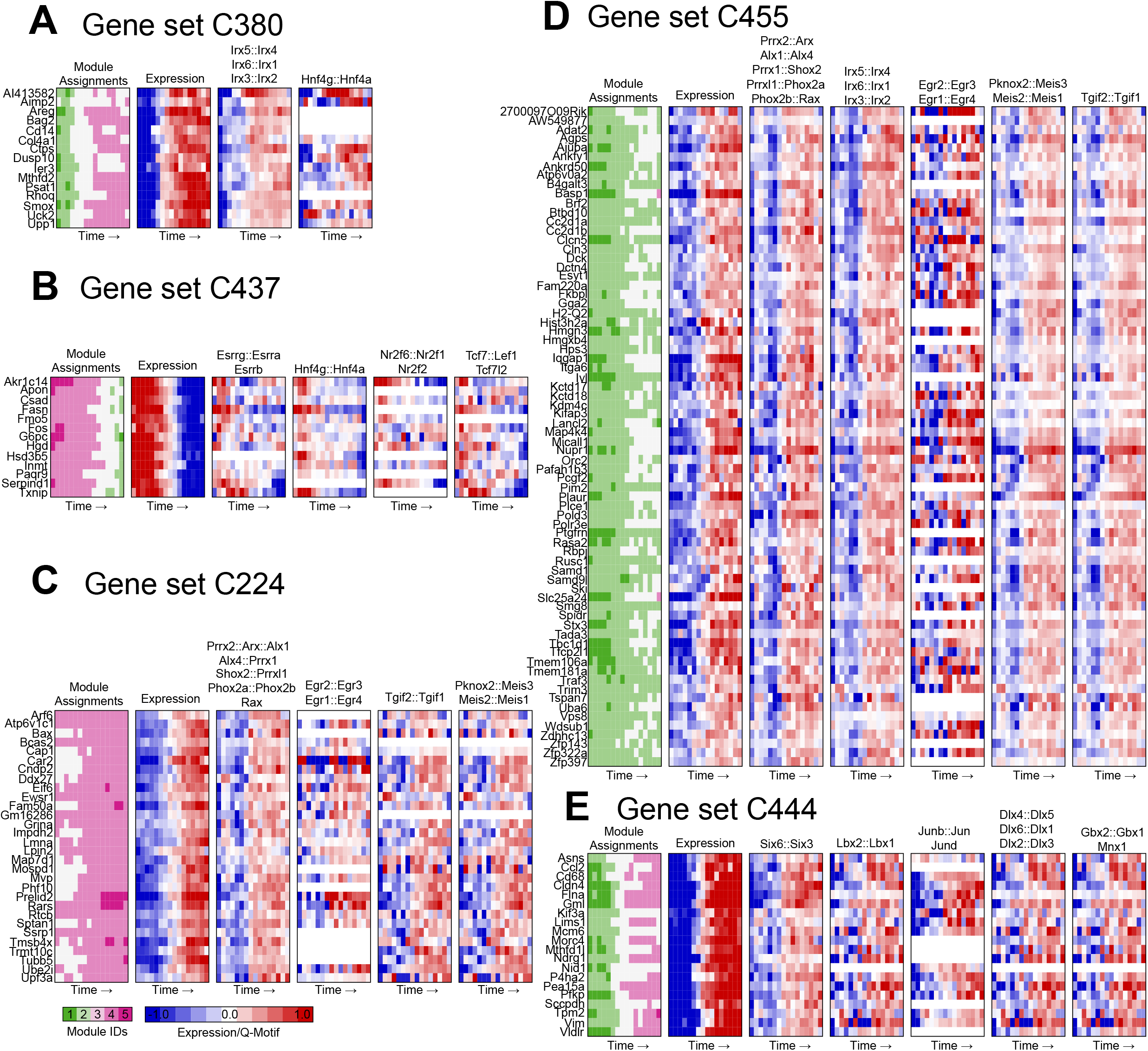
Selected transitioning gene sets in the hepatocyte dedifferentiation dataset. Each panel shows the member genes of a transitioning gene set (label on top). The columns show the module assignment of each gene, followed by its expression level in each cell type (Exp). The subsequent groups of columns are the levels of the regulator (Q-motif value) on the gene promoters. The name of the regulator is specified at the top of each block of columns. All significant regulators are shown.

### DRMN application on early embryonic lineage specification

To demonstrate the utility of DRMN on hierarchical trajectories, we considered a dataset profiling early differentiation of human embryonic stem cells (H1) into four lineages, mesendoderm, mesenchymal, neural progenitors, trophoblast [13]. In addition to accessibility, this dataset measured eight different histone marks including H3K4me1, H3K4me2, H3K4me3, H3K27ac, H3K9ac, H3K79me2, H3K36me3 and H3K27me3. We applied DRMN to this dataset and identified modules representing five major levels of expression, 1-5, with 1 representing the lowest expression and 5 the highest (**Figure 8A**). The extent of gene conservation in modules depended on the module with modules 1 and 2 exhibiting low conservation, while modules 3 and 4 exhibiting high conservation. The low conservation of modules 1 and 2 is consistent with our observations in the reprogramming study (**Figure 8B**). Gene Ontology (GO) process enrichment showed that the repressed module is enriched for developmental and lineage specific functions, while the induced modules tended to be enriched for cell cycle and translation related processes (**Additional file 2**). The most repressed module (Module 1) exhibited the largest extent of cell type specific enrichments, while the induced module (Module 5) was enriched for similar processes. As such the GO enrichment was not able to capture lineage-specific processes in the induced modules.

**Figure 8.**
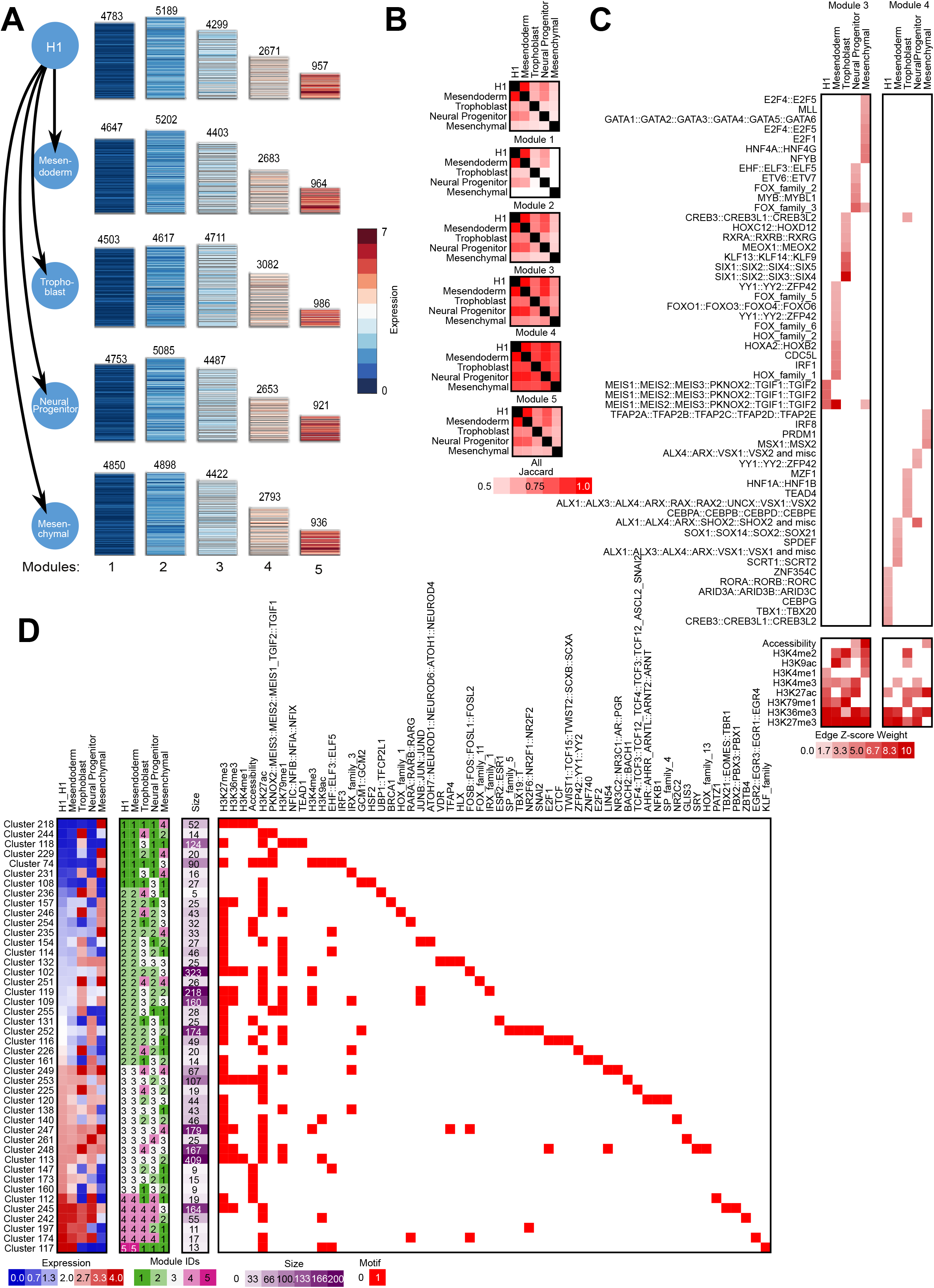
Application of DRMNs to the hepatocyte dedifferentiation dataset using accessibility and Q-motifs for the feature set. **A**. Show are gene expression patterns of *k* = 5 modules for each lineage (major row). The number above the heatmap is the number of genes in that module. **B**. Similarity of modules across lineages as measured by F-score. The color intensity is proportional to the match. **C**. Inferred regulators for each modules across time. Only modules with highest expression and that had TFs as regulators (3 and 4) are shown. The red intensity is proportional the z-score significance of the regression weight of a regulator **D**. Transitioning gene sets exhibiting changes into the high expression modules (3, 4, 5). Shown are the mean expression levels of genes in the gene set (left, red-blue heat map), the module assignment (second), the number of genes in each module (third) and the set of regulators for each gene set (white-red heat map).

We next examined the *cis*-regulatory elements selected by DRMN, focusing on the two highly expressed modules, 3 and 4 (Module 5 was associated only with histone marks). Similar to the reprogramming study, histone mark associations were more conserved across cell types than TFs, with the elongation mark, H3K36me3 and repressive mark, H3K27me3 being among the most conserved marks predicted across cell types. Among the TFs associated with each module, several have known lineage-specific roles (**Figure 8C**). For example, we found several neuronal lineage regulators including FOX [40] and MYB [41] proteins in module 3 of the Neural Progenitor cell type, and VSX1 [42] and SHOX2 [43] in Mesendoderm module 4. Similarly, TEAD4, a regulator important for trophoblast self renewal [44] was associated with Trophoblast Module 4.

Finally, we examined the transitioning gene sets and identified a total of 6988 (of 17904) genes that changed their module state. We grouped these into 122 gene sets of at least 5 genes and included 6730 genes (**Additional file 3**). We focused on gene sets transitioning into the high modules of 3, 4 and 5 and predicted regulators for them (**Methods**) and identified a total of 44 (**Figure 8D**). We found several gene sets with lineage-specific patterns of expression. For example, some gene sets showed lineage specific up-regulation for Mesenchymal (C218, C235, **Figure 9A, B**), neuronal (C226, C154, **Figure 9C, D**) and the trophoblast lineages (C131, C161 **Figure 9E, F**). Several of the gene sets involved known lineagespecific genes, e.g., NEUROD1 for the neuronal lineage-specific gene set C154 [45] (**Figure 9D**), as well as regulators important for multiple lineage (e.g., E2F1 and E2F2 TFs in C161 [46], **Figure 9F**). Several gene sets were up-regulated in multiple lineages (C138 and C112, **Figure 9G, H**), which could indicate regulatory programs with multi-potent potential. Taken together, these results provided insight into the interplay of chromatin marks, accessibility and TFs to specify lineage specific expression patterns.

**Figure 9.**
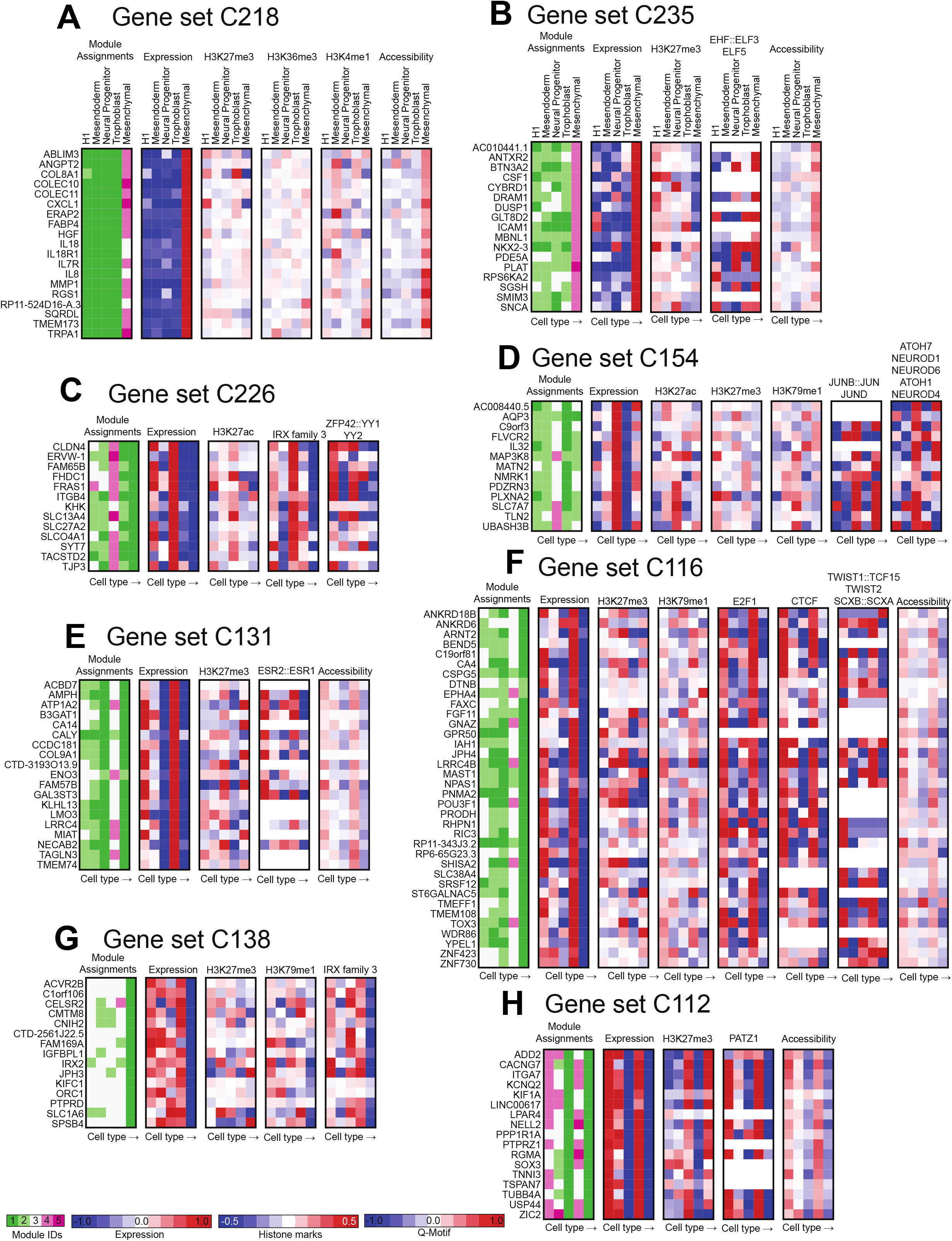
Selected transitioning gene sets in the ESC differentiation dataset. Each panel shows the member genes of a transitioning gene set (label on top). The columns show the module assignment of each gene, followed by its expression level in each cell type (Exp). The subsequent groups of columns are the levels of the regulator (Histone or Q-motif value) on the gene promoters. The name of the regulator is specified at the top of each block of columns. All significant regulators are shown. **A-B**. Mesenchymal specific gene sets. **C-D**. Neural progenitor specific lineage. **E-F**. Gene sets with trophoblast-specific expression. **G-H**. Gene sets exhibiting multi-lineage expression.

## Discussion

To gain insight into regulatory network dynamics associated with cell type specific expression patterns, time course and hierarchically related datasets measuring transcriptomes and epigenomes of a dynamic process are becoming increasingly available [5–7, 13, 47]. Analyzing these datasets to identify the underlying gene regulatory network dynamics that drive context-specific expression changes is a major challenge. This is because of the large number of variables measured in each context (time point or cell type), but low sample size for each context. In this work, we developed DRMN, that simplifies genome scale regulatory networks from individual genes to gene modules and infers regulatory program for each module in all the input conditions. Using DRMN, one can characterize the major transcriptional patterns during a dynamic process and identify transcription factors and epigenomic signals that are responsible for these transitions.

Central to DRMN’s modeling framework is to jointly learn the regulatory programs for each cell type or time point by using multi-task learning. Using two different approaches to multi-task learning, we show that joint learning of regulatory programs is advantageous compared to a simpler approach of learning regulatory programs independently per condition. Furthermore, predictive modeling of expression that also clusters genes into expression groups is more powerful than simple clustering. Such models have improved generalizability and are able to capture fine-grained expression variation as a function of the upstream regulatory state of a gene.

DRMN offers a flexible framework to integrate a variety of regulatory genomic signals. In its simplest form, DRMN can be applied to expression datasets with sequence-specific motifs to learn a predictive regulatory model. In its more general form, DRMN can integrate other types of regulatory signals such as genome-wide chromatin accessibility, histone modification and transcription factor profiles measured using sequencing and array technologies. Furthermore DRMN is applicable to datasets of different experimental designs such as short time series (e.g., the reprogramming study), long time series (e.g., the hepatocyte dedifferentiation study) and hierarchically related cell types on lineages (e.g. in the cellular differentiation study).

We used DRMN’s predictive modeling framework to systematically study the utility of different celltype specific measurements such as chromatin marks and accessibility to predict expression. When combining ATAC-seq with chromatin marks to predict expression, we did not see a substantial improvement in predictive power. This is likely because of the large number of chromatin marks in our dataset. However, in both array and sequencing data we found that combining sequence features (motifs) and histone modifications had the highest predictive power. In our application to the Chronis et al dataset, we did not observe a substantial advantage of using accessible motif instances (Q-motif) as opposed to all motif instances. This is most likely due to the sparsity of the feature space of Q-motifs which came from the gene promoter. A direction of future work would be to incorporate ATAC-seq signal from more distal regions by using genome-wide chromosome conformation capture assays [48–50]. Another direction would be to use more generic sequence features, such as k-mers [51] to enable the discovery of novel regulatory elements and offer great flexibility in capturing sequence specificity and its role in predictive models of expression.

We applied DRMNs to distinct types of dynamic processes which involved cell fate transitions: mouse reprogramming from a differentiated fibroblast cell state to a pluripotent state (3-4 cell types), hepatocyte dedifferentiation (16 time points), and forward differentiation of human embryonic stem cells to different lineages (5 cell types). DRMN application identified the major patterns of expression in these datasets as well as dynamic, transitioning genes that changed their expression state over time or condition. Interestingly, when comparing across processes we found that the repressed modules were least conserved in the reprogramming and differentiation study, while in the de-differentiation study the repressed module was more conserved. Accordingly, biological processes were repressed in a cell state specific manner in the reprogramming and forward differentiation experiment, while we saw a broad down-regulation of developmental processes in the hepatocyte dedifferentiation dataset. Furthermore, there were greater changes in expression in the reprogramming time course compared to dedifferentiation indicative of the different dynamics in the two processes. Importantly DRMN was able to identify key regulators for gene modules as well as for the transitioning genes that included both novel and known regulators for each process. For example, DRMN identified several ESC specific regulators in the reprogramming dataset, liver transcription factors such as HNF4G::HNF4A in the dedifferentiation dataset, and lineage-specific factors in the differentiation dataset. These predictions offer testable hypothesis of regulators driving important expression dynamics in cell fate transitions.

## Conclusion

Cell type-specific gene expression patterns are established by a complex interplay of multiple regulatory levels including transcription factor binding, genome accessibility and histone modifications. DRMN offers a powerful and flexible framework to define cell type specific gene regulatory programs from time series and hierarchically related regulatory genomic datasets. As additional datasets that profile epigenomic and transcriptomic dynamics of specific processes become available, methods like DRMN will become increasingly useful to examine regulatory network dynamics underlying context-specific expression.

## Materials and methods

### Dynamic Regulatory Module Networks (DRMN)

### DRMN model description

DRMN is based on a module-based representation of a regulatory network, where we group genes into modules and learn a regulatory program for each gene module. This module-based representation is more appropriate when the number of samples per condition are too few to perform conventional gene regulatory network inference of estimating the regulators of individual genes for each time point [14, 52]. The regulatory program for the modules of any one condition is referred to as Regulatory Module Network (RMN). DRMN infers regulatory module networks for multiple biological conditions (e.g., cell types, time points) related by a time course or a lineage tree, where each condition has a small number of samples (e.g, one or two) but several types of measurements, such as RNA-seq, ChIP-seq and ATAC-seq. For ease of description, we will assume we have a set of related cell types, however the same description applies to multiple time points or related conditions.

DRMN takes as inputs: (i) **X**, an *N × C* matrix of cell type-specific expression values for *N* genes in *C* cell types, assuming we have a single measurement of a gene in each cell type; (ii) *Y* = {**Y**_1_, *…*, **Y**_*C*_} collection of feature matrices, one for each cell type, *c*. Each **Y**_*c*_ is an *N × F* matrix, with the *i*^th^ row specifying the values of *F* features for gene *i*; and (iii) a lineage tree, *τ* which describes how the *C* cell types are related, (iv) *k* the number of gene modules. The DRMN model is defined by a set of RMNs, **R** = {**R**_1_, *…*, **R**_*C*_} linked via the lineage tree *τ*, and transition probability distributions **Π** = {Π_1_, …, Π_*C*_} (**Figure 1**). For each cell type *c*, **R**_*c*_ =*< G*_*c*_, Θ_*c*_ *>*, where *G*_*c*_ is graph defining the set of edges between *F* features and *k* modules, and Θ_*c*_, are the parameters of regression functions (one for each module) that relate the selected regulatory features to the expression of the genes in a module. Transition matrices {Π_1_, …, Π_*C*_} capture the dynamics of the module assignments in one cell type *c* given its parent cell type in the lineage tree *τ*. Specifically, Π_*c*_(*i, j*) is the probability of any gene being in module *j* given that its parental assignment is to module *i*. For the root cell type, this is simply a prior probability over modules. The values inside each feature matrix, **Y**_*c*_ can be either context-independent (*e*.*g*., a sequence-based motif network) or context-specific (*e*.*g*., a motif network informed by accessibility value, histone modifications). Given the above inputs, DRMN optimization aims to optimize the posterior likelihood of the model given the data, *P* (**R**|**X, Y**_**1**_, *…*, **Y**_*C*_). Based on Bayes rule, this can be rewritten as:

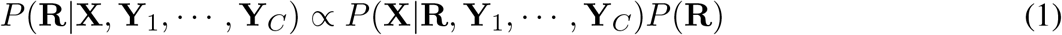

Here *P* (**X**|**R, Y**_1_, *…*, **Y**_*C*_) is the data likelihood given the regulatory program. For the second term, we assume that *P* (**R**|**Y**_1_, *…*, **Y**_*C*_) = *P* (**R**). The data likelihood can be decomposed over the individual cell types as: *P* (**X**|**R, Y**) = Π _*c*_ *P* (**X**_*c*_|**R**_*c*_, **Y**_*c*_). Within each cell type, this is modeled using a mixture of predictive models, one model for each module. For the prior, *P* (**R**), we used two formulations to enable sharing information: DRMN-Structure Prior (DRMN-ST) defines a structure prior over the graph structures *P* (*G*_1_, …, *G*_*C*_) while DRMN-FUSED uses a regularized regression framework and implicitly defines priors on the *P* (Θ_1_, *…*, Θ_*C*_). In both frameworks, we share information between the cell types/conditions to learn the regulatory programs of each cell type/condition.

#### DRMN-ST: Structure prior approach

In DRMN-ST, information is shared by specifying a structure prior, *P* (*G*_1_, …, *G*_*C*_), defined only over the graphs. The parameters are set to their maximum likelihood setting. *P* (*G*_1_, …, *G*_*C*_) determines how information is shared between different cell types at the level of the network structure and encourages similarity of features, indicative of regulators, between cell types. *P* (*G*_1_, …, *G*_*C*_) is computed using the transition matrices {Π_1_, …, Π_*C*_} and decomposes over individual regulator-module edges within each cell type as follows:

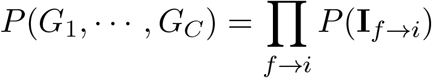

where

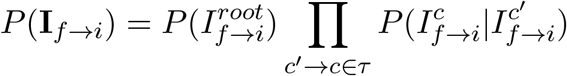

where 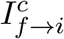 is an indicator function for the presence of the edge *f* → *i* for cell type *c*, between a regulator *f* and a module *i*. To define 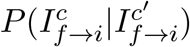, we use the transition probability of the modules as:

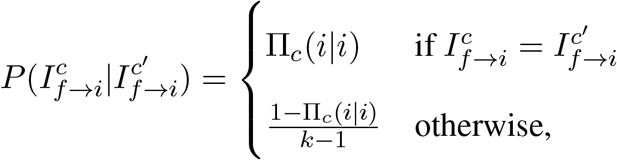

where *k* is the number of modules. Here the first option gives the probability of maintaining the same state from parent to child cell types (is present or absent in both *c* and *c ′*), and the second option gives the probability of changing the edge state. We incorporated the structure prior within a greedy hill climbing algorithm for estimating the DRMN model (see Section **DRMN learning** for more details of the learning algorithm).

#### DRMN-FUSED: Parameter prior approach

In DRMN-FUSED, we use a fused group LASSO formulation to share information between the cell types by defining the following objective:

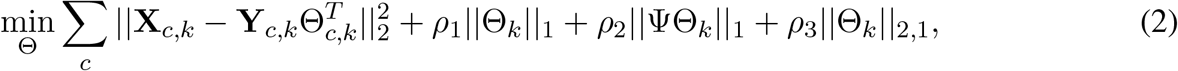

where **X**_*c,k*_ is the expression vector of genes in module *k* in cell line *c*, **Y**_*c,k*_ is the feature matrix for genes in module *k*, and Θ_*c,k*_ is a 1 *× F* vector of regression coefficients for the same module and cell line (non-zero values correspond to selected features). Θ_*k*_ is the *C* by *F* matrix resulting from stacking up the Θ_*c,k*_ vectors as rows. Ψ is a *C* − 1 by *C* matrix, encoding the lineage tree. Each row of Ψ corresponds to a branch of the tree and each column corresponds to a cell type. If row *i* of Ψ corresponds to a branch *c ′* → *c* in the lineage tree, Ψ(*i, c*_1_) = 1, Ψ(*i, c*_2_) = −1, and all other values in that row are 0. ΨΘ_*k*_ is a *C* − 1 by *F*, where row *i* correspond to Θ_*c′,k*_ − Θ_*c,k*_, the difference between regression coefficients of cell lines *c ′* and *c*. ||.||_1_ denotes *l*_1_-norm (sum of absolute values), ||.||_2_ denotes *l*_2_-norm (square root of sum of square of value), and ||.||_2,1_ denotes *l*_2,1_-norm (sum of *l*_2_-norm of columns of the given matrix). *ρ*_1_, *ρ*_2_ and *ρ*_3_ correspond to hyper parameters, with *ρ*_1_ for sparsity penalty, *ρ*_2_ to enforce similarity between selected features of consecutive cell lines in the lineage tree, and *ρ*_3_ to enforce selecting the same features for all cell types. Thus, *ρ*_2_ controls the extent to which more closely related cell types are closer in their regression weights, which *ρ*_3_ controls the extent to which all the cell types share similarity in their regulatory programs. We set these hyper parameters based on cross-validation by performing a grid search for *ρ*_1_, *ρ*_2_ and *ρ*_3_ as described in the dataset-specific application sections. We implemented the optimization algorithm by extending the algorithm described in the MALSAR Matlab package [53] to handle branching topologies. The learning algorithm uses an accelerated gradient method [54, 55] to minimize the objective function above.

#### DRMN learning

DRMNs are learned by optimizing the DRMN score (**Eqn** 1), using an Expectation Maximization (EM) style algorithm that searches over the space of possible graphs for a local optimum (**Algorithm 1**). In the Maximization (M) step, we estimate transition parameters (M1 step) and the regulatory program structure (M2 step). In the Expectation (E) step, we compute the expected probability of a gene’s expression profile to be generated by one of the regulatory programs. The M2 step uses multi-task learning to jointly learn the regulatory programs for all cell types using either the framework of DRMN-ST or DRMN-FUSED.

##### Algorithm 1: DRMN Algorithm

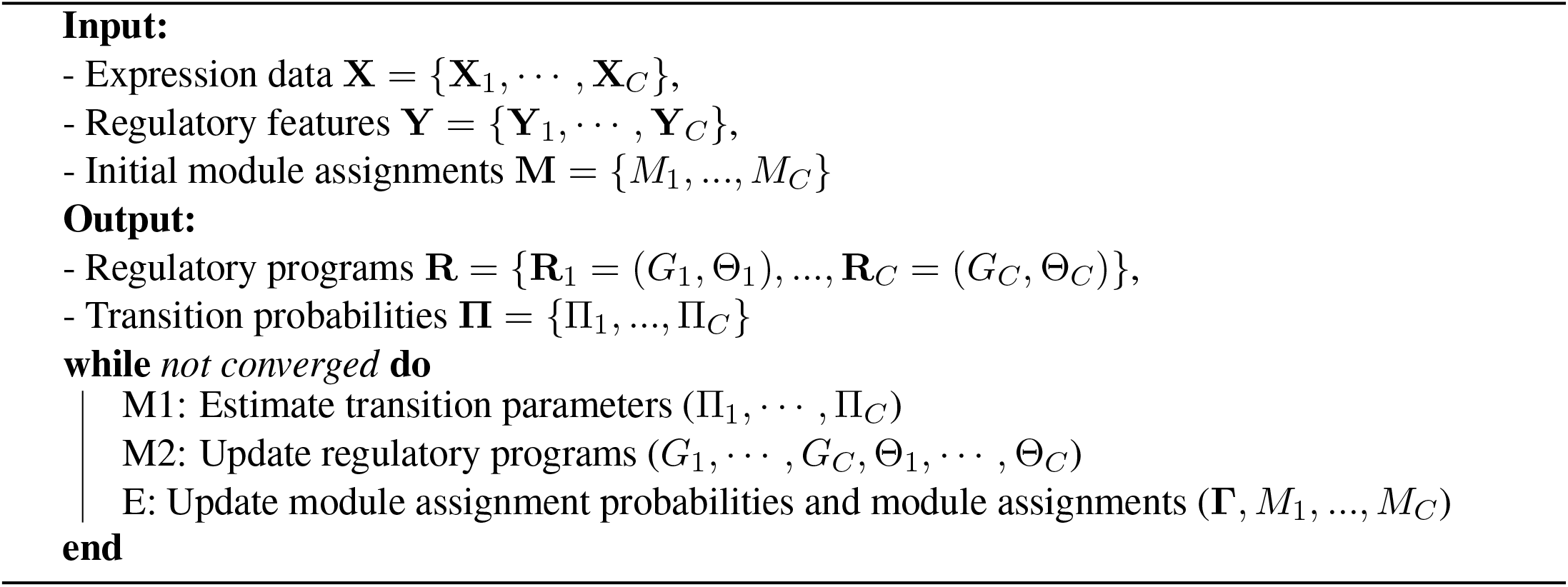

#### Estimate transition parameters (M1 step)

Let 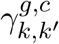 be the probability of gene *g* in cell type *c* to belong to module *k*, and, in its parent cell type *c ′*, to module *k ′*. We calculate the probability of transitioning from *k ′* in *c ′* to *k* in *c* as 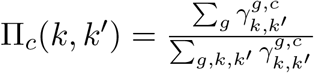.

#### Update regulatory programs (M2 step)

Recall that the regulatory program for each cell type *c* is **R**_*c*_ =*< G*_*c*_, Θ_*c*_ *>*, where *G*_*c*_ is a set of regulatory interactions *f* → *i* from a regulatory feature *f* to a module *i*, and Θ_*c*_ are the parameters of a regression function for each module that relates the selected regulatory features to the expression of the genes in a module. We assume that the expression levels are generated by a mixture of experts, each expert corresponding to a module. Each expert uses a multivariate normal distribution, with mean *µ*_*c,i*_ and covariance Σ_*c,i*_. Let *V*_*c,i*_ denote the set of regulatory features in *G*_*c,i*_. *µ*_*c,i*_ is |*V*_*c,i*_| + 1 *×* 1 and Σ_*c,i*_ is square matrix with |*V*_*c,i*_| + 1 rows and columns, where the additional dimension is for expression. Given, *µ*_*c,i*_ and Σ_*c,i*_, let *θ*_*c,i*_ denote the conditional mean of the expression variable given all the regulatory features and let 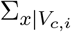 be the conditional variance. Let **X**_*c*_(*g*) denote the expression level of a gene *g* in cell type *c*. The probability of **X**_*c*_(*g*) from module *i* in cell type *c* is:

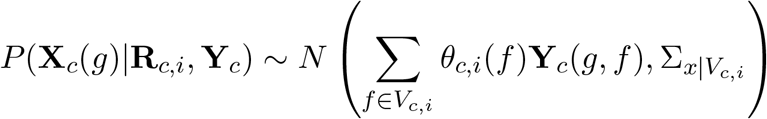

The estimation of these multivariate distributions is done one module at a time across all cell types, however the procedure is specific to DRMN-ST and DRMN-FUSED. In the DRMN-ST approach, the regulatory interactions are learned one module, across all cell types at a time using a greedy hill-climbing framework. At initialization, for each module *i, G*_*c,i*_ is an empty graph, and the Gaussian parameters are computed as the empirical mean and variance of the genes initially assigned to module *i* in each cell type. In each iteration, we score each potential regulatory feature based on its improvement to the likelihood of the model, and choose the regulator with maximum improvement. This regulator is added to the module’s regulatory program for all cell types for which it improves the cell type-specific likelihood. In DRMN-FUSED, the structure and parameters of **R**_*c,i*_ are learned by optimizing the objective in **Eqn 2** using an accelerated gradient method [54, 55]. See **Additional file 1 Supplemental text** for more details.

#### Update soft module assignments (E step)

Let 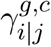 be the probability of gene *g* in cell type *c* to belong to module *i*, given that in its parent cell type *c ′*, it belonged to module *j*. We also introduce 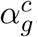, a vector of size *k ×* 1 where each element 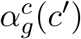 specifies the probability of observations given the parent state is *c ′*. We estimate the probabilities using a dynamic programming procedure, where values at internal nodes in the lineage tree are computed using the values for all descendent nodes, down to the leaves.

If *c* is a leaf node, we calculate

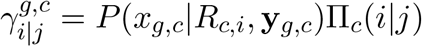

where the first term is the probability of observing expression of gene *g* in cell type *c* in module *i* (given its regulatory program and regulatory features), and the second is the probability of transitioning from module *j* in the parent cell type *c ′* to module *i* in cell type *c*.

For a non-leaf cell type *c*:

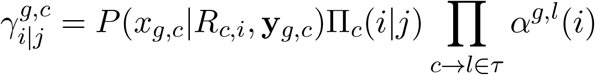

For both internal and leaf cell types, we write the joint probability of the (*j, i*) pair as 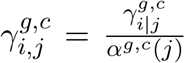, where 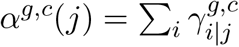 is the probability of *g*’s expression in any module given parent module *j*.

#### Termination

DRMN inference runs for a set number of iterations or until convergence. Final module assignments are computed as maximum likelihood assignments using a dynamic programming approach. While module assignments between consecutive iterations do not change significantly, the final module assignments are significantly different from the initial module assignments, and predictive power of model significantly improves as iterations progress (though improvements are small after 10 iterations, **Additional file 1 Figure S7**). In our experiments, we ran DRMN for up to ten iterations. When using greedy hill climbing approach, the M2 step was run until up to five regulators were added per module.

### Feature sets tested in DRMN

We initially examined DRMN with a variety of feature sets to study the utility of different feature combinations for predicting expression. Most of the feature set types were examined with the reprogramming array and sequencing datasets. We considered the following features for each gene to predict its expression.

#### Motif

We defined motif features based on the presence of a motif instance of a transcription factor (TF) within the gene’s promoter region, defined as *±*2500 around the gene TSS. We downloaded a meta-compilation of position weight matrices (PWMs) from various resources (see dataset-specific sections for details) for human (e.g., CisBP) or mouse (CisBP and Sherwood et al [56]). We applied FIMO [57] to scan the mouse or human genome for significant motif instances (*p <* 1*e* − 5). For each gene, we generated a vector of motif presence, one dimension for each motif with the value equal to the −log_10_(*p*-value) of a motif instance. If a gene had multiple motif instances for the same motif, we used the most significant instance (smallest *p*-value).

#### Histone

For datasets with histone modifications measured, we used each histone mark as a separate feature. This included 8 features for the reprogramming array dataset, 9 features for the sequencing dataset and 8 features for the H1ESC differentiation dataset. The feature value was the aggregated count value around the gene TSS followed by log transformation.

#### Histone + Motif

This was the concatenation of histone modification features (Histone) where available, with the motif features for each gene.

#### Accessibility

The Accessibility feature was a single feature repressing the aggregated ATAC-seq or DNase-seq reads around the gene promoter. After aggregating to the gene promoter, we quantile normalized and log transformed the values.

#### Accessibility + Motif

This feature set represents the concatenation of the ATAC feature with the Motif feature set for each gene.

#### Q-Motif

This feature set represents sequence-specific motif features scored by the ATAC-seq/DNase-seq signal producing a total of as many features as there are motifs with significant instances. We used BedTools (`bedtools genomecov -ibam input.bam -bg -pc > output.counts`) to obtain the aggregated signal on each base pair. We defined the feature value as the log-transformed mean read count under each motif instance. If multiple instances of the same motif were mapped to the same transcript, the signal was summed. If a TF was mapped to multiple transcripts of the same gene, or multiple motifs of the same TF were mapped to the same gene, the max value was used.

#### Histone + Accessibility + Motif

This feature set represents the concatenation of the Histone feature set, ATAC feature and the Motif feature set.

#### Histone + Q-Motif

Similar to the Histone + Motif feature set, Histone + Q-Motif represents the concatenation of the Histone and Q-Motif feature set for each gene.

#### Histone + Accessibility + Q-Motif

This feature set is similar to Histone+ATAC+Motif and represents the concatenation of Histone and Q-Motif feature sets with the ATAC feature.

### Application of DRMN to different datasets

We applied DRMN to study regulatory network dynamics to four different processes (a) A microarray time course dataset of cellular reprogramming from mouse embryonic fibroblasts (MEFs) to pluripotent cells, (b) A sequencing time course dataset of cellular reprogramming using the same system as (a), (c) A sequencing time course dataset profiling dedifferentation of hepatocyte cells, (d) differentiation of H1ESC cells to different developmental lineages. Below we describe the dataset processing, feature generation, hyper-parameter selection and analysis of transitioning gene sets.

#### Reprogramming array data

This dataset had measurements of gene expression and eight chromatin marks in three cell types: mouse embryonic fibroblasts (MEFs), partially reprogrammed induced pluripotent stem cells (pre-iPSCs), and induced pluripotent stem cells (iPSCs), collected from multiple publications [12, 58–60]. The expression values of 15,982 genes was measured by microarray. Eight chromatin marks were measured by ChIP-on-chip (chromatin immunoprecipitation followed by promoter microarray). For each gene promoter, each mark’s value was averaged across a 8000-bp region associated with the promoter. The chromatin marks included those associated with active transcription (H3K4me3, H3K9ac, H3K14ac, and H3K18ac), repression (H3K9me2, H3K9me3, H3K27me3), and transcription elongation (H3K79me2).

For this dataset, we considered the following set of features: Motif, Histone, Histone+Motif. For the Motif feature, we used the motif collection available with the PIQ software [56] from http://piq.csail.mit.edu/, which were sourced from multiple databases [61–63]. From the full motif list, we used only those annotated as transcription factor proteins [64]. This resulted in a total of 353 TFs. We used FIMO to find motif instances using the mm9 mouse genome. Motif instances for the same TF were further aggregated into a single feature per gene by selecting the motif instance with the most significant *p*-value. This resulted in a total of 353 dimensions for the Motif feature.

We used the array and the reprogramming dataset to study the effect of hyper-parameters on the performance of DRMN-FUSED with different feature sets. The hyper-parameters are *ρ*_1_ (sparsity in each task), *ρ*_2_ (selection of more similar features for closely related cell types) and *ρ*_3_ (selection of similar features for all cell types). We performed a grid search on a wide range of parameters values:

*ρ*_1_ *∈* {0.5, 1, 2, 5, 10, 20, 30, 40, 50, 60, 70, 80, 90, 100, 110, 120, 130}, *ρ*_2_ *∈* {0, 10, 20, 30, 40, 50} and *ρ*_3_ = {0, 10, 20, 30, 40, 50} (**Additional file 1 Figure S8**). We used a three-fold cross validation setting and computed average correlation between predicted and true expression for each module and cell line. We used the average over modules and cell lines to assess DRMN performance for particular feature set and hyper-parameter setting. We observe that increase in *ρ*_3_ was generally not beneficial for the Motif feature for all *k*, number of modules, and for *k ≥* 7 when using Histone and Histone+Motif (blue *ρ*_3_=0) *vs*. cyan, *ρ*_3_=50, **Additional file 1 Figure S8**). For a fixed value of *ρ*_3_, increasing sparsity *ρ*_1_ is beneficial for the Histone and Histone + Motif features, upto *ρ*_1_ = 30 − 60, beyond which the performance decreases or does not improve. The *ρ*_2_ feature was also most useful when using the Histone feature.

Final results of DRMN were obtained by using the Motif+Histone feature with DRMN-FUSED. We inspected the performance across different hyper-parameter settings using these features and found the top 5 configurations to be: {(50, 50, 30), (50, 10, 30), (90, 10, 0), (40, 40, 50), (30, 50, 50)}. Inspection of the heat maps DRMN results showed that the results were similar across these configurations, where we selected the first configuration (50, 50, 30) for DRMN application on the full dataset. DRMN modules were interpreted using Gene Ontology enrichment using a Hyper-geometric T-test. Transitioning gene sets were defined based on genes that change

#### Reprogramming sequencing data

This dataset generated by Chronis et al. [7], assayed gene expression with RNA-seq, nine chromatin marks with ChIP-seq, and chromatin accessibility with ATAC-seq in different stages of reprogramming (MEF, MEF48 (48 hours after start of the reprogramming process), pre-iPSC, and embryonic stem cells (ESC)). The chromatin marks included H3K27ac, H3K27me3, H3K36me3, H3K4me1, H3K4me2, H3K4me3, H3K79me2, H3K9ac, and H3K9me3. We aligned all sequencing reads to the mouse mm9 reference genome using Bowtie2 [65]. For RNA-seq data, we quantified expression to TPMs using RSEM [66] and applied a log transform. After removing unexpressed genes (TPM*<*1), we had 17,358 genes. For the ChIP-seq and ATAC-seq datasets, we obtained per-base pair read coverage using BEDTools [67], aggregated counts within *±*2, 500 bps of a gene’s transcription start site and applied log transformation.

For this dataset, we considered all the feature types described in the section, **Feature sets tested in DRMN**. The Motif feature was generated in a similar manner as in the reprogramming array dataset. In addition, we included, Q-Motif, Accessibility and their combinations with the Histone feature.

Similar to the array dataset, we used this dataset to study the effect of hyper-parameters, *ρ*_1_, *ρ*_2_ and *ρ*_3_ on the performance of DRMN-FUSED using a similar three-fold cross-validation framework (**Additional file 1 Figures S9, S10, S11**). Similar to the array dataset, increasing values of *ρ*_3_ was not beneficial for Motif or Q-Motif alone. Higher value of *ρ*_3_ was useful for some of settings of histones features combined with Accessibility, Motif or Q-motif (*k* = 3, 5 for generally lower values of *ρ*_1_, **Additional file 1 Figures S9, S10, S11**). We next investigate the impact of *ρ*_1_ and *ρ*_2_, for different values of *ρ*_3_. We observe that when using histone features (Histone+Motif, Histone+Accessibility+Motif, Histone+Accessibility+Q-Motif,), increase in *ρ*_2_ (increasing the similarity of inferred networks) improves the predictive power of the method. Increase in *ρ*_1_ is beneficial for these features upto a limit (typically, *ρ*_1_=60 or 70). Conversely, for the feature sets that do not use histone features (Motif, Q-Motif, and Accessibility+Motif), increase in *ρ*_1_ (sparser models) decrease the predictive power of the model, which is consistent with the decrease in performance of Motif features in the array dataset.

Final results from DRMN for this dataset are obtained by applying DRMN-FUSED to the Histone+Accessibility+Q-Motif feature set with *k* = 7 modules in three-fold cross-validation mode and examining the top 5 configurations. For this dataset, these were {(10,50,40), (20,50,40), (10,50,10), (40,50,0), (10,50,0)}. Next we inspected the heat maps of DRMNs and determined that the results are largely similar. We present results for one of the configurations:*ρ*_1_ = 10, *ρ*_2_ = 50, *ρ*_3_ = 40. Similar to the reprogramming array dataset we tested for GO enrichment and defined transitioning gene sets. We next used a simple regression model to predict regulators for each gene set (See **Section**, “Identification of transitioning gene sets and their regulators”).

#### Hepatocyte dedifferentiation time course data

The dedifferentiation time course consisted of samples were extracted from adult mouse liver and gene expression and chromatin accessibility were assayed by RNA-seq and ATAC-seq, respectively, at 0 hrs, 0.5 hrs, 1 hrs, 2 hrs, 4 hrs, 6 hrs, 8 hrs, 10 hrs, 12 hrs, 14 hrs, 16 hrs, 18 hrs, 20 hrs, 22 hrs, 24 hours, and 36 hrs (16 time points in total, [26]). All sequencing reads were aligned to mouse mm10 reference genome using Bowtie2 [65], and gene expression was quantified using RSEM [66]. Any gene with TPM= 0 in all time points was removed, resulting in 14,794 genes with measurement in at least one time point. Per-base pair read coverage for ATAC-seq was obtained using BEDTools [67], and counts were aggregated within *±*2, 500 bps of a gene’s transcription start site. Both gene expression and accessibility data were quantile normalized across 16 time points and then log transformed.

DRMN was applied with Accessibility and Q-Motif as the feature set with *k* = 5 modules using the DRMN-FUSED implementation. Q-Motif features were generated similar to the reprogramming dataset. We used a utility in the PIQ package [56] to identify motif instances in *±*2, 500bp of a gene TSS using PWMs from CIS-BP database [68]. Motif instances were scored by the ATAC-seq signal resulting in a total of 2,856 features. The Q-Motif Features were quantile normalized and log transformed across the 16 time points.

To determine the appropriate settings for the hyper-parameters for DRMN-FUSED, we scanned the following range of values of values: *ρ*_1_ *∈* {30, 50, 70, 90, 110, 130, 150}, *ρ*_2_ *∈* {0, 10, 30, 50, 70} and *ρ*_3_ = {0, 10, 30, 50, 70} within a three-fold cross-validation setting (**Additional file 1 Figure S12**) and picked the best setting based on overall prediction error (as described in the procedure for the reprogramming dataset). We used the Accessibility+Q-Motif feature set and *k* = 5 for the number of modules. We found the top 5 parameter configurations to be {(70,10,70), (70,0,70), (90,0,50), (70,30,50), (70,50,30)}. The modules from these five configurations looked similar. We finally did a full run of DRMN-FUSED with (*ρ*_1_ = 70, *ρ*_2_ = 10, *ρ*_3_ = 70), which had the best prediction error. The DRMN modules were interpreted with GO enrichment analysis. We defined transitioning gene sets from the DRMN modules and used a multi-task group LASSO regression approach to select regulators for each gene set (See **Section**, “Identification of transitioning gene sets and their regulators” for more details).

#### H1ESC differentiation into different lineages

The differentiation dataset from Xie et al. [13] profiled gene expression, accessibility and histone modifications in human embryonic stem cells (hESCs) and four lineages derived from hESCs, mesendoderm, neural progenitor, trophoblast-like, and mesenchymal stem cells. Gene expression was measured with RNA-seq, histone modifications were measured with ChIP-seq and chromatin accessibility with DNaseseq. The dataset included eight histone modification marks: H3K27ac, H3K27me3, H3K36me3, H3K4me1, H3K4me2, H3K4me3, H3K79me1, and H3K9ac. We aligned all sequencing reads to the human hg19 reference genome using Bowtie2 [65]. For RNA-seq data, we quantified expression to TPMs using RSEM [66], quantile normalized across cell lines and applied a log transform, resulting in 17,899 genes with expression across all cell lines. For both the ChIP-seq (chromatin marks) and DNase-seq data, we obtained per-base pair read coverage using BEDTools [67] and aggregated counts within *±*2, 500 bps of a gene transcription start site (TSS).

DRMN was applied using the full set of features, Accessibility, Q-Motif, Histone with *k* = 5 modules. Q-Motif features were obtained using Cis-BP human motif PWM collection by applying utility tools from the PIQ software package [56] on *±*2500bp of a gene TSS. Each motif instance was scored with the DNAseseq signal resulting in a total of 2,998 features. These features were quantile normalized and log transformed across the cell lines.

To determine the appropriate settings of the hyper-parameters, we performed three fold cross-validation with hyper parameter values in the following range: *ρ*_1_ *∈* {30, 50, 70, 90, 110, 130, 150}, *ρ*_2_ *∈* {0, 10, 30, 50, 70} and *ρ*_3_ *∈* {0, 10, 30, 50, 70} (**Additional file 1 Figure S13**). The top 5 configurations were {(150,0,30), (30,0,0), (150,0,10), (30,0,10), (130,0,30)}. The results from these settings were similar. The results reported here were generated by applying DRMN on the full dataset with *ρ*_1_ = 150, *ρ*_2_ = 0, *ρ*_3_ = 30. We note that for this dataset, as the relationships between the cell types is captured by a two-level tree, the *ρ*_2_ parameter likely does not have a large effect and most of the task sharing is sufficiently captured by the global *ρ*_3_ parameter. Once modules were defined we analyzed them as described in the above sections and generated transitioning gene sets. We identified regulators for the transitioning gene sets using our simple regression-based approach (See **Section**, “Identification of transitioning gene sets and their regulators” for more details).

### Identification of significant cis-regulatory elements and regulatory features per module

To analyse the inferred regulatory programs in DRMN models, we sought to identify regulatory features that are significantly different across the contexts. Briefly, for each regulatory feature selected for each module, we calculated the z-score of inferred DRMN edge weights across all conditions (cell-type/time point). In each module features with a significant edge (z-score*>* 1 for reprogramming datasets, and z-score*>* 3 for dedifferentiation and ES differentiation datasets) in at least one conditon were selected. The complete list of identified features are available in **Additional file 5**.

### Identification of transitioning gene sets and their regulators

A transitioning gene is defined as a gene for which its module assignment changes in at least one time point/cell line. We grouped the genes into transitioning gene sets using a hierarchical clustering approach with city block as a distance metric and 0.05 as a distance threshold. We developed two strategies for selecting regulators for transitioning gene sets: Simple linear regression for short time courses (with 5 or fewer time points) and Multi-Task Group LASSO (MTG-LASSO) that is suitable for longer time courses. Both approaches take as input a list of genes from a transitioning set and predict the regulators that are likely responsible for the change in expression over time.

#### Simple linear regression approach

To identify regulators associated with transitioning gene sets in datasets with ≤ 5 samples (the two reprogramming datasets and H1ESC differentiation dataset), we used a simple regression-based approach, that sought to find regulators that can explain the overall variation in expression of genes in the set. Briefly, for each transitioning gene set with *n* genes across *C* cell types, we created a *n*.*C ×* 1 expression vector by stacking of the *C*-dimensional vector for each of the genes across the *C* cell types. We repeated this procedure for all the features associated with the gene set to produce a *n*.*C × F* matrix, where *F* is the total number of features in our dataset. Next, we used regularized regression to select which features are most predictive of the expression levels. Any regularized regression framework can be used; we used the sparsity imposing regression framework of the MERLIN algorithm [69], which uses a probabilistic framework with a prior term (tuned using a hyper-parameter) to infer sparser models. We ran MERLIN (with default settings) on each transitioning gene set to identify regulatory features associated with that set. We finally filtered the predicted regulators per gene set by assessing the correlation of the regulator/feature with the gene expression of a gene across the cell lines and included a regulator if was correlated to at least 5 genes with a Pearson’s correlation of 0.6 or higher.

#### Multi-Task Group Lasso

To identify regulators associated with transitioning gene sets in datasets with *≥* 6 samples, e.g., the dedifferentiation dataset, we used a multi-task regression framework called Multi-Task Group Lasso (MTG-Lasso). In MTG-Lasso, we perform multiple regression tasks, one for each gene in the set to select regulatory features as predictors for each gene’s expression levels. The “group” penalty of MTG-Lasso enables us to select the same regulator for all genes in the gene set but with different parameters. The regulatory feature corresponds to a “group” with the regression weights for the regulator across all genes to be the in the same group. MTG-Lasso selects or unselects entire groups of regression weights. The MTG-Lasso objective for each gene set *s* is:

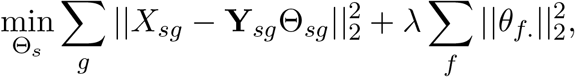

Here *X*_*sg*_ is the *C ×* 1 vector of expression values over *C* samples for gene *g*, and **Y**_*sg*_ is the *C × F* matrix of regulatory features for *g* over samples. Θ_*sg*_ = [*θ*_1*g*_, *…, θ*_*F g*_]^T^ is the *F ×* 1 vector of regression weights for predicting *g*’s expression from the *F* regulatory features. The first term denotes the regression task for each gene *g*, while the second term denotes the L1/L2 regularization required for the MTG-LASSO framework. The sum over *f* imposes the L1 penalty selecting a small number of groups (one *θ*_*f*._ for each feature *f*), and the 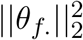 imposes the L2 norm for smoothness of the regression coefficients across genes. *λ* is the hyper-parameter controlling for the strength of the regularization.

We applied MTG-Lasso to each transitioning gene set using the MATLAB SLEP v4.1 package [70] to infer the most predictive regulatory features for the gene set. We performed leave-one-out cross-validation, where one sample is left out from training, a model is fit on the remaining samples and used to predict the left out sample. We computed a confidence for each feature based on the percentage of models in which the feature is selected. Additionally, we computed a *p*-value for the selection of each regulator by comparing the number of times it was selected to a null distribution of feature selection obtained from randomizing the data 40 times and training MTG-Lasso models. A feature was selected as a regulator if it was in at least 60% of the trained values and had a *p*-value*<* 0.05. We tried different hyper-parameter values (*λ ∈* {0.1, 0.2, 0.3, 0.4, 0.5, 0.6, 0.7, 0.8, 0.9, 0.99}) and selected the value that resulted in reasonable number of regulators across all transitioning gene sets (*λ* = 0.7). Once regulators were selected for each gene set, we further filtered the features based on the pearson correlation of the gene expression and feature values as in the simple regression case.

## Availability

The DRMN code is available at https://github.com/Roy-lab/drmn along with usage instructions. Data pre-processing and feature generation scripts are available at https://github.com/Roy-lab/drmn_utils. DRMN outputs have been provided as **Additional file 6** (the necessary mapping between motif names and TF names are available in **Additional file 7**).

## Acknowledgment

This work was made possible in part by NIH NIGMS grant 1R01GM117339 to S.R. The authors thank the Center for High Throughput Computing (CHTC) for computing resources.

## Additional files

**Additional File 1**. Supplemental data including supplementary text and **Figures S1**-**S13**.

**Additional File 2**. GO terms enriched in each module of each cell line or time point. Each sheet correspond to one dataset.

**Additional File 3**. List of genes that change module across cell lines/time points. Each sheet correspond to one data set.

**Additional File 4**. Regulators associated with the transitioning gene sets. The file contains three zip files (for reprogramming dataset from from Chronis et al., the dedifferentiation dataset, and the ESC differentiation dataset). Each zip file contains a file per transitioning gene set, where first column is the name of regulator, second column is the name of target (Exp for all files), and the last column correspond to edge weight (regression coefficient for linear regression and confidence for MTG LASSO).

**Additional File 5**. Significant interaction per module, where significance was defined as z-score higher than the given threshold (1 for reprogramming datasets and 3 for the other two datasets). Each sheet corresponds to one dataset.

**Additional File 6**. The zip file contains four zip files (one for each dataset) containing the DRMN outputs of inferred networks and module assignments per cell line/time point.

**Additional File 7**. The zip file contains map of motif names to gene names for human and mouse CIS-BP motifs. It also contains the shortened names used in different figures.

## References

[1] Alvaro J. Gonzalez, Manu Setty, and Christina S. Leslie. Early enhancer establishment and regulatory locus complexity shape transcriptional programs in hematopoietic differentiation. Nature genetics, 47(11):1249–1259, Nov 2015. 26390058[pmid].

[2] Hatice U. Osmanbeyoglu, Fumiko Shimizu, Angela Rynne-Vidal, Petar Jelinic, Samuel C. Mok, Gabriela Chiosis, Douglas A. Levine, and Christina S. Leslie. Chromatin-informed inference of transcriptional programs in gynecologic and basal breast cancers. bioRxiv, 2018.

[3] Richard A. Young. Control of the embryonic stem cell state. Cell, 144(6):940–954, 2011.

[4] Tong Ihn Lee and Richard A Young. Transcriptional regulation and its misregulation in disease. Cell, 152(6):1237–1251, Mar 2013.

[5] Joseph A Wamstad, Jeffrey M Alexander, Rebecca M Truty, Avanti Shrikumar, Fugen Li, Kirsten E Eilertson, Huiming Ding, John N Wylie, Alexander R Pico, John A Capra, Genevieve Erwin, Steven J Kattman, Gordon M Keller, Deepak Srivastava, Stuart S Levine, Katherine S Pollard, Alisha K Holloway, Laurie A Boyer, and Benoit G Bruneau. Dynamic and coordinated epigenetic regulation of developmental transitions in the cardiac lineage. Cell, 151:206–220, September 2012.

[6] David Lara-Astiaso, Assaf Weiner, Erika Lorenzo-Vivas, Irina Zaretsky, Diego Adhemar Jaitin, Eyal David, Hadas Keren-Shaul, Alexander Mildner, Deborah Winter, Steffen Jung, Nir Friedman, and Ido Amit. Chromatin state dynamics during blood formation. Science (New York, N.Y.), 345:943–949, August 2014.

[7] Constantinos Chronis, Petko Fiziev, Bernadett Papp, Stefan Butz, Giancarlo Bonora, Shan Sabri, Jason Ernst, and Kathrin Plath. Cooperative binding of transcription factors orchestrates reprogramming. Cell, 168:442–459.e20, January 2017.

[8] Xianjun Dong, Melissa C Greven, Anshul Kundaje, Sarah Djebali, James B Brown, Chao Cheng, Thomas R Gingeras, Mark Gerstein, Roderic Guigó, Ewan Birney, and Zhiping Weng. Modeling gene expression using chromatin features in various cellular contexts. Genome biology, 13:R53, June 2012.

[9] Thais G do Rego, Helge G Roider, Francisco A T de Carvalho, and Ivan G Costa. Inferring epigenetic and transcriptional regulation during blood cell development with a mixture of sparse linear models. Bioinformatics (Oxford, England), 28:2297–2303, September 2012.

[10] Jason Ernst, Oded Vainas, Christopher T Harbison, Itamar Simon, and Ziv Bar-Joseph. Reconstructing dynamic regulatory maps. Molecular systems biology, 3:74, 2007.

[11] Wuming Gong, Naoko Koyano-Nakagawa, Tongbin Li, and Daniel J Garry. Inferring dynamic gene regulatory networks in cardiac differentiation through the integration of multi-dimensional data. BMC bioinformatics, 16:74, March 2015.

[12] Sushmita Roy and Rupa Sridharan. Chromatin module inference on cellular trajectories identifies key transition points and poised epigenetic states in diverse developmental processes. Genome research, 27:1250–1262, July 2017.

[13] Wei Xie, Matthew D. Schultz, Ryan Lister, Zhonggang Hou, Nisha Rajagopal, Pradipta Ray, John W. Whitaker, Shulan Tian, R. David Hawkins, Danny Leung, Hongbo Yang, Tao Wang, Ah Young Y. Lee, Scott A. Swanson, Jiuchun Zhang, Yun Zhu, Audrey Kim, Joseph R. Nery, Mark A. Urich, Samantha Kuan, Chia-an A. Yen, Sarit Klugman, Pengzhi Yu, Kran Suknuntha, Nicholas E. Propson, Huaming Chen, Lee E. Edsall, Ulrich Wagner, Yan Li, Zhen Ye, Ashwinikumar Kulkarni, Zhenyu Xuan, Wen-Yu Y. Chung, Neil C. Chi, Jessica E. Antosiewicz-Bourget, Igor Slukvin, Ron Stewart, Michael Q. Zhang, Wei Wang, James A. Thomson, Joseph R. Ecker, and Bing Ren. Epigenomic analysis of multilineage differentiation of human embryonic stem cells. Cell, 153(5):1134–1148, May 2013.

[14] Eran Segal, Michael Shapira, Aviv Regev, Dana Pe’er, David Botstein, Daphne Koller, and Nir Friedman. Module networks: identifying regulatory modules and their condition-specific regulators from gene expression data. Nat. Genet., 34(2):166–176, May 2003.

[15] Su-In Lee, Aimée M. Dudley, David Drubin, Pamela A. Silver, Nevan J. Krogan, Dana Pe’er, and Daphne Koller. Learning a prior on regulatory potential from eQTL data. PLoS Genet, 5(1):e1000358.#x002B;, January 2009.

[16] Vladimir Jojic, Tal Shay, Katelyn Sylvia, Or Zuk, Xin Sun, Joonsoo Kang, Aviv Regev, Daphne Koller, Immunological Genome Project Consortium, Adam J. Best, Jamie Knell, Ananda Goldrath, Vladimir Joic, Daphne Koller, Tal Shay, Aviv Regev, Nadia Cohen, Patrick Brennan, Michael Brenner, Francis Kim, Tata Nageswara Rao, Amy Wagers, Tracy Heng, Jeffrey Ericson, Katherine Rothamel, Adriana Ortiz-Lopez, Diane Mathis, Christophe Benoist, Natalie A. Bezman, Joseph C. Sun, Gundula Min-Oo, Charlie C. Kim, Lewis L. Lanier, Jennifer Miller, Brian Brown, Miriam Merad, Emmanuel L. Gautier, Claudia Jakubzick, Gwendalyn J. Randolph, Paul Monach, David A. Blair, Michael L. Dustin, Susan A. Shinton, Richard R. Hardy, David Laidlaw, Jim Collins, Roi Gazit, Derrick J. Rossi, Nidhi Malhotra, Katelyn Sylvia, Joonsoo Kang, Taras Kreslavsky, Anne Fletcher, Kutlu Elpek, Angelique Bellemarte-Pelletier, Deepali Malhotra, and Shannon Turley. Identification of transcriptional regulators in the mouse immune system. Nat Immunol, 14(6):633–643, Jun 2013.

[17] Xiaozhen Dai, Xiaoqing Yan, Kupper A. Wintergerst, Lu Cai, Bradley B. Keller, and Yi Tan. Nrf2: Redox and metabolic regulator of stem cell state and function. Trends in Molecular Medicine, 26(2):185–200, Feb 2020.

[18] N. Miyashita, M. Horie, H. I. Suzuki, M. Saito, Y. Mikami, K. Okuda, R. C. Boucher, M. Suzukawa, A. Hebisawa, A. Saito, and T. Nagase. FOXL1 Regulates Lung Fibroblast Function via Multiple Mechanisms. Am J Respir Cell Mol Biol, 63(6):831–842, 12 2020.

[19] Cristina Tavera-Montañez, Sarah J. Hainer, Daniella Cangussu, Shellaina J. V. Gordon, Yao Xiao, Pablo Reyes-Gutierrez, Anthony N. Imbalzano, Juan G. Navea, Thomas G. Fazzio, and Teresita Padilla-Benavides. The classic metal-sensing transcription factor mtf1 promotes myogenesis in response to copper. FASEB journal : official publication of the Federation of American Societies for Experimental Biology, 33(12):14556–14574, Dec 2019. 31690123[pmid].

[20] Pedro Vallecillo-García, Mickael Orgeur, Sophie vom Hofe-Schneider, Jürgen Stumm, Verena Kappert, Daniel M. Ibrahim, Stefan T. Börno, Shinichiro Hayashi, Frédéric Relaix, Katrin Hildebrandt, Gerhard Sengle, Manuel Koch, Bernd Timmermann, Giovanna Marazzi, David A. Sassoon, Delphine Duprez, and Sigmar Stricker. Odd skipped-related 1 identifies a population of embryonic fibroadipogenic progenitors regulating myogenesis during limb development. Nature Communications, 8(1):1218, Oct 2017.

[21] Quan M. Phan, Gracelyn M. Fine, Lucia Salz, Gerardo G. Herrera, Ben Wildman, Iwona M. Driskell, and Ryan R. Driskell. Lef1 expression in fibroblasts maintains developmental potential in adult skin to regenerate wounds. eLife, 9:e60066, Sep 2020. 32990218[pmid].

[22] T. Ikeda, J. Zhang, T. Chano, A. Mabuchi, A. Fukuda, H. Kawaguchi, K. Nakamura, and S. Ikegawa. Identification and characterization of the human long form of Sox5 (L-SOX5) gene. Gene, 298(1):59–68, Sep 2002.

[23] Takashi Ikeda, Takafusa Hikichi, Hisashi Miura, Hirofumi Shibata, Kanae Mitsunaga, Yosuke Yamada, Knut Woltjen, Kei Miyamoto, Ichiro Hiratani, Yasuhiro Yamada, Akitsu Hotta, Takuya Yamamoto, Keisuke Okita, and Shinji Masui. Srf destabilizes cellular identity by suppressing cell-type-specific gene expression programs. Nature Communications, 9(1):1387, Apr 2018.

[24] Elo Madissoon, Anastasios Damdimopoulos, Shintaro Katayama, Kaarel Krjutškov, Elisabet Einarsdottir, Katariina Mamia, Bert De Groef, Outi Hovatta, Juha Kere, and Pauliina Damdimopoulou. Pleomorphic adenoma gene 1 is needed for timely zygotic genome activation and early embryo development. Scientific Reports, 9(1):8411, Jun 2019.

[25] Greetje Elaut, Tom Henkens, Peggy Papeleu, Sarah Snykers, Mathieu Vinken, Tamara Vanhaecke, and Vera Rogiers. Molecular mechanisms underlying the dedifferentiation process of isolated hepatocytes and their cultures. Current Drug Metabolism, 7(6):629–660, 2006.

[26] Morten Seirup, Srikumar Sengupta, Scott Swanson, Brian E. McIntosh, Mike Colins, Li-Fang Chu, Zhang Cheng, David U. Gorkin, Bret Duffin, Jennifer M. Bolin, Cara Argus, Ron Stewart, and James A. Thomson. Rapid changes in chromatin structure during dedifferentiation of primary hepatocytes in vitro. bioRxiv, 2020.

[27] Tom Luedde and Robert F. Schwabe. Nf-κb in the liver–linking injury, fibrosis and hepatocellular carcinoma. Nature reviews. Gastroenterology & hepatology, 8(2):108–118, Feb 2011. 21293511[pmid].

[28] Masaji Sakaguchi, Weikang Cai, Chih-Hao Wang, Carly T. Cederquist, Marcos Damasio, Erica P. Homan, Thiago Batista, Alfred K. Ramirez, Manoj K. Gupta, Martin Steger, Nicolai J. Wewer Albrechtsen, Shailendra Kumar Singh, Eiichi Araki, Matthias Mann, Sven Enerbäck, and C. Ronald Kahn. Foxk1 and foxk2 in insulin regulation of cellular and mitochondrial metabolism. Nature Communications, 10(1):1582, Apr 2019.

[29] J. Le Lay and K. H. Kaestner. The Fox genes in the liver: from organogenesis to functional integration. Physiol Rev, 90(1):1–22, Jan 2010.

[30] F. Clotman, P. Jacquemin, N. Plumb-Rudewiez, C. E. Pierreux, P. Van der Smissen, H. C. Dietz, P. J. Courtoy, G. G. Rousseau, and F. P. Lemaigre. Control of liver cell fate decision by a gradient of TGF beta signaling modulated by Onecut transcription factors. Genes Dev, 19(16):1849–1854, Aug 2005.

[31] Ilaria Laudadio, Isabelle Manfroid, Younes Achouri, Dominic Schmidt, Michael D. Wilson, Sabine Cordi, Lieven Thorrez, Laurent Knoops, Patrick Jacquemin, Frans Schuit, Christophe E. Pierreux, Duncan T. Odom, Bernard Peers, and Frederic P. Lemaigre. A feedback loop between the liver-enriched transcription factor network and mir-122 controls hepatocyte differentiation. Gastroenterology, 142(1):119–129, Jan 2012.

[32] Ewa Wandzioch, A°sa Kolterud, Maria Jacobsson, Scott L. Friedman, and Leif Carlsson. Lhx2–/–mice develop liver fibrosis. Proceedings of the National Academy of Sciences, 101(47):16549–16554, 2004.

[33] Megan F. Cole, Sarah E. Johnstone, Jamie J. Newman, Michael H. Kagey, and Richard A. Young. Tcf3 is an integral component of the core regulatory circuitry of embryonic stem cells. Genes & development, 22(6):746–755, Mar 2008. 18347094[pmid].

[34] Masayuki Uemura, E. Scott Swenson, Marianna D. A. Gaça, Frank J. Giordano, Michael Reiss, and Rebecca G. Wells. Smad2 and smad3 play different roles in rat hepatic stellate cell function and alpha-smooth muscle actin organization. Molecular biology of the cell, 16(9):4214–4224, Sep 2005. 15987742[pmid].

[35] Wenjie Yu, Xiao Li, Steven Eliason, Miguel Romero-Bustillos, Ryan J. Ries, Huojun Cao, and Brad A. Amendt. Irx1 regulates dental outer enamel epithelial and lung alveolar type ii epithelial differentiation. Developmental biology, 429(1):44–55, Sep 2017. 28746823[pmid].

[36] Jiangying Liu, You Wang, Morris J. Birnbaum, and Doris A. Stoffers. Three-amino-acid-loop-extension homeodomain factor meis3 regulates cell survival via pdk1. Proceedings of the National Academy of Sciences, 107(47):20494–20499, 2010.

[37] Jian Ming Khor, Jennifer Guerrero-Santoro, and Charles A. Ettensohn. Genome-wide identification of binding sites and gene targets of alx1, a pivotal regulator of echinoderm skeletogenesis. Development, 146(16), 2019.

[38] Nancy Magee and Yuxia Zhang. Role of early growth response 1 in liver metabolism and liver cancer. Hepatoma research, 3:268–277, 2017. 29607419[pmid].

[39] Swetha Rudraiah, Xi Zhang, and Li Wang. Nuclear receptors as therapeutic targets in liver disease: Are we there yet? Annual Review of Pharmacology and Toxicology, 56(1):605–626, 2016. PMID: 26738480.

[40] Anna L. M. Ferri, Wei Lin, Yannis E. Mavromatakis, Julie C. Wang, Hiroshi Sasaki, Jeffrey A. Whitsett, and Siew-Lan Ang. Foxa1 and foxa2 regulate multiple phases of midbrain dopaminergic neuron development in a dosage-dependent manner. Development, 134(15):2761–2769, 2007.

[41] J. Malaterre, T. Mantamadiotis, S. Dworkin, S. Lightowler, Q. Yang, M. I. Ransome, A. M. Turnley, N. R. Nichols, N. R. Emambokus, J. Frampton, and R. G. Ramsay. c-Myb is required for neural progenitor cell proliferation and maintenance of the neural stem cell niche in adult brain. Stem Cells, 26(1):173–181, Jan 2008.

[42] Cédric Francius, María Hidalgo-Figueroa, Stéphanie Debrulle, Barbara Pelosi, Vincent Rucchin, Kara Ronellenfitch, Elena Panayiotou, Neoklis Makrides, Kamana Misra, Audrey Harris, Hessameh Hassani, Olivier Schakman, Carlos Parras, Mengqing Xiang, Stavros Malas, Robert L. Chow, and Frédéric Clotman. Vsx1 transiently defines an early intermediate v2 interneuron precursor compartment in the mouse developing spinal cord. Frontiers in molecular neuroscience, 9:145–145, Dec 2016. 28082864[pmid].

[43] A. Scott, H. Hasegawa, K. Sakurai, A. Yaron, J. Cobb, and F. Wang. Transcription factor short stature homeobox 2 is required for proper development of tropomyosin-related kinase B-expressing mechanosensory neurons. J Neurosci, 31(18):6741–6749, May 2011.

[44] Biswarup Saha, Avishek Ganguly, Pratik Home, Bhaswati Bhattacharya, Soma Ray, Ananya Ghosh, M. A. Karim Rumi, Courtney Marsh, Valerie A. French, Sumedha Gunewardena, and Soumen Paul. Tead4 ensures postimplantation development by promoting trophoblast self-renewal: An implication in early human pregnancy loss. Proceedings of the National Academy of Sciences, 117(30):17864–17875, 2020.

[45] Abhijeet Pataskar, Johannes Jung, Pawel Smialowski, Florian Noack, Federico Calegari, Tobias Straub, and Vijay K Tiwari. Neurod1 reprograms chromatin and transcription factor landscapes to induce the neuronal program. The EMBO Journal, 35(1):24–45, 2016.

[46] M. M. Ouseph, J. Li, H. Z. Chen, T. Pecot, P. Wenzel, J. C. Thompson, G. Comstock, V. Chokshi, M. Byrne, B. Forde, J. L. Chong, K. Huang, R. Machiraju, A. de Bruin, and G. Leone. Atypical E2F repressors and activators coordinate placental development. Dev Cell, 22(4):849–862, Apr 2012.

[47] Daria Bunina, Nade Abazova, Nichole Diaz, Kyung-Min Noh, Jeroen Krijgsveld, and Judith B. Zaugg. Genomic rewiring of sox2 chromatin interaction network during differentiation of escs to postmitotic neurons. Cell Systems, 10(6):480–494.e8, Jun 2020.

[48] James Fraser, Iain Williamson, Wendy A. Bickmore, and Josée Dostie. An overview of genome organization and how we got there: from fish to hi-c. Microbiology and Molecular Biology Reviews, 79(3):347–372, 2015.

[49] Elzo de Wit and Wouter de Laat. A decade of 3c technologies: insights into nuclear organization. Genes & Development, 26(1):11–24, 2012.

[50] David U. Gorkin, Danny Leung, and Bing Ren. The 3d genome in transcriptional regulation and pluripotency. Cell Stem Cell, 14(6):762–775, Jun 2014.

[51] Manu Setty and Christina S. Leslie. Seqgl identifies context-dependent binding signals in genomewide regulatory element maps. PLOS Computational Biology, 11(5):1–21, 05 2015.

[52] ANSHUL Kundaje, STEVE Lianoglou, Xuejing LI, DAVID Quigley, MARTA Arias, CHRIS H. Wiggins, LI Zhang, and CHRISTINA Leslie. Learning regulatory programs that accurately predict differential expression with medusa. Annals of the New York Academy of Sciences, 1115(1):178–202, 2007.

[53] Jiayu Zhou, Jianhui Chen, and Jieping Ye. Malsar: Multi-task learning via structural regularization – user’s manual version 1.1, 2012.

[54] Yu. Nesterov. Smooth minimization of non-smooth functions. Mathematical Programming, 103(1):127–152, May 2005.

[55] Yu. Nesterov. Gradient methods for minimizing composite objective function. CORE Discussion Papers 2007076, Université catholique de Louvain, Center for Operations Research and Econometrics (CORE), 2007.

[56] Richard I. Sherwood, Tatsunori Hashimoto, Charles W. O’Donnell, Sophia Lewis, Amira A. Barkal, John Peter van Hoff, Vivek Karun, Tommi Jaakkola, and David K. Gifford. Discovery of directional and nondirectional pioneer transcription factors by modeling dnase profile magnitude and shape. Nat Biotechnol, 32(2):171–178, Feb 2014.

[57] Charles E. Grant, Timothy L. Bailey, and William Stafford Noble. Fimo: scanning for occurrences of a given motif. Bioinformatics, 27(7):1017–1018, Apr 2011.

[58] Nimet Maherali, Rupa Sridharan, Wei Xie, Jochen Utikal, Sarah Eminli, Katrin Arnold, Matthias Stadtfeld, Robin Yachechko, Jason Tchieu, Rudolf Jaenisch, Kathrin Plath, and Konrad Hochedlinger. Directly reprogrammed fibroblasts show global epigenetic remodeling and widespread tissue contribution. Cell stem cell, 1:55–70, June 2007.

[59] Rupa Sridharan, Jason Tchieu, Mike J Mason, Robin Yachechko, Edward Kuoy, Steve Horvath, Qing Zhou, and Kathrin Plath. Role of the murine reprogramming factors in the induction of pluripotency. Cell, 136:364–377, January 2009.

[60] Rupa Sridharan, Michelle Gonzales-Cope, Constantinos Chronis, Giancarlo Bonora, Robin McKee, Chengyang Huang, Sanjeet Patel, David Lopez, Nilamadhab Mishra, Matteo Pellegrini, Michael Carey, Benjamin A Garcia, and Kathrin Plath. Proteomic and genomic approaches reveal critical functions of H3K9 methylation and heterochromatin protein-1γ in reprogramming to pluripotency. Nature cell biology, 15:872–882, July 2013.

[61] V Matys, E Fricke, R Geffers, E Gössling, M Haubrock, R Hehl, K Hornischer, D Karas, A E Kel, O V Kel-Margoulis, D-U Kloos, S Land, B Lewicki-Potapov, H Michael, R Münch, I Reuter, S Rotert, H Saxel, M Scheer, S Thiele, and E Wingender. Transfac: transcriptional regulation, from patterns to profiles. Nucleic acids research, 31:374–378, January 2003.

[62] Albin Sandelin, Wynand Alkema, Pär Engström, Wyeth W Wasserman, and Boris Lenhard. Jaspar: an open-access database for eukaryotic transcription factor binding profiles. Nucleic acids research, 32:D91–D94, January 2004.

[63] Michael F Berger, Anthony A Philippakis, Aaron M Qureshi, Fangxue S He, Preston W Estep, and Martha L Bulyk. Compact, universal dna microarrays to comprehensively determine transcriptionfactor binding site specificities. Nature biotechnology, 24:1429–1435, November 2006.

[64] Timothy Ravasi, Harukazu Suzuki, Carlo V. Cannistraci, Shintaro Katayama, Vladimir B. Bajic, Kai Tan, Altuna Akalin, Sebastian Schmeier, Mutsumi Kanamori-Katayama, Nicolas Bertin, Piero Carninci, Carsten O. Daub, Alistair R. R. Forrest, Julian Gough, Sean Grimmond, Jung-Hoon Han, Takehiro Hashimoto, Winston Hide, Oliver Hofmann, Atanas Kamburov, Mandeep Kaur, Hideya Kawaji, Atsutaka Kubosaki, Timo Lassmann, Erik van Nimwegen, Cameron R. MacPherson, Chihiro Ogawa, Aleksandar Radovanovic, Ariel Schwartz, Rohan D. Teasdale, Jesper Tegnér, Boris Lenhard, Sarah A. Teichmann, Takahiro Arakawa, Noriko Ninomiya, Kayoko Murakami, Michihira Tagami, Shiro Fukuda, Kengo Imamura, Chikatoshi Kai, Ryoko Ishihara, Yayoi Kitazume, Jun Kawai, David A. Hume, Trey Ideker, and Yoshihide Hayashizaki. An atlas of combinatorial transcriptional regulation in mouse and man. Cell, 140(5):744–752, March 2010.

[65] Ben Langmead and Steven L Salzberg. Fast gapped-read alignment with bowtie 2. Nature methods, 9:357–359, March 2012.

[66] Bo Li and Colin N Dewey. Rsem: accurate transcript quantification from rna-seq data with or without a reference genome. BMC bioinformatics, 12:323, August 2011.

[67] Aaron R. Quinlan and Ira M. Hall. BEDTools: a flexible suite of utilities for comparing genomic features. Bioinformatics, 26(6):841–842, 01 2010.

[68] Matthew T. Weirauch, Ally Yang, Mihai Albu, Atina G. Cote, Alejandro Montenegro-Montero, Philipp Drewe, Hamed S. Najafabadi, Samuel A. Lambert, Ishminder Mann, Kate Cook, Hong Zheng, Alejandra Goity, Harm van Bakel, Jean-Claude Lozano, Mary Galli, Mathew G. Lewsey, Eryong Huang, Tuhin Mukherjee, Xiaoting Chen, John S. Reece-Hoyes, Sridhar Govindarajan, Gad Shaulsky, Albertha J M. Walhout, François-Yves Bouget, Gunnar Ratsch, Luis F. Larrondo, Joseph R. Ecker, and Timothy R. Hughes. Determination and inference of eukaryotic transcription factor sequence specificity. Cell, 158(6):1431–1443, Sep 2014.

[69] Sushmita Roy, Stephen Lagree, Zhonggang Hou, James A. Thomson, Ron Stewart, and Audrey P. Gasch. Integrated module and Gene-Specific regulatory inference implicates upstream signaling networks. PLoS Comput. Biol., 9(10):e1003252.+, October 2013.

[70] Jolanta Jura, Paulina Wegrzyn, Michal Korostynski, Krzysztof Guzik, Malgorzata Oczko-Wojciechowska, Michal Jarzab, Malgorzata Kowalska, Marcin Piechota, Ryszard Przewlocki, and Aleksander Koj. Identification of interleukin-1 and interleukin-6-responsive genes in human monocyte-derived macrophages using microarrays. Biochimica et Biophysica Acta (BBA) - Gene Regulatory Mechanisms, 1779(6):383–389, 2008.

