## Supplementary Materials for "Dynamic regulatory module networks for inference of cell type-specific transcriptional networks"

immediate

#### Contents

**1 Supplementary Methods** **2**

1.1 DRMN-FUSED . . . . . . . . . . 2

1.2 DRMN-ST . . . . . . . . . . 7

**2 Supplementary Figures** **9**

**References** **36**

### 1 Supplementary Methods

We implemented two versions of DRMN for sharing information across tasks, where each task is a cell line: DRMN-FUSED and DRMN-ST. Below we describe the details of these algorithms.

#### 1.1 DRMN-FUSED

As described in the main text, DRMN-FUSED uses the fused Group LASSO formulation to share information between the cell types per module. The objective of DRMN-FUSED is:

$$\min_{\Theta} \sum_{c} ||\mathbf{Y}_{c,k} - \mathbf{X}_{c,k}\Theta_{c,k}^{\intercal}||_2^2 + \rho_1||\Theta_k||_1 + \rho_2||\Psi\Theta_k||_1 + \rho_3||\Theta_k||_{2,1}, \tag{1}$$

where $\mathbf{Y}_{c,k}$ is the $n_k \times 1$ expression vector of genes in module $k$ in cell line $c$, $\mathbf{X}_{c,k}$ is the $n_k \times F$ feature matrix for genes in module $k$ in cell line $c$, and $\Theta_{c,k}$ is a $1 \times F$ vector of regression coefficients for the same module and cell line. $\Theta_k$ is the $C$ by $F$ matrix resulting from stacking up the $\Theta_{c,k}$ vectors as rows. $\Psi$ is a $C - 1$ by $C$ matrix, encoding the lineage tree. Each row correpond to an edge in the tree, with values 1 and $-1$ for the nodes that the edge connects, meaning that $||\Psi\Theta_k||_1$ is equivalent of $\sum_{c_1 \to c_2 \in \Psi} ||\Theta_{c_1,k} - \Theta_{c_2,k}||_1$. $\rho_1, \rho_2$ and $\rho_3$ correspond to hyper parameters, with $\rho_1$ for sparsity penalty, $\rho_2$ to enforce similarity between selected features as specified by $\Psi$, and $\rho_3$ to enforce selecting the same features for all cell types.

The FUSED LASSO objective is not smooth and therefore requires special handling, that is they have a smooth ($l_2$ loss) and a non-smooth part (Fused LASSO part). We follow the implementation in the MALSAR package [1], which makes use of Nesterov's accelerated gradient method (AGM) for composite functions [2, 3]. The MALSAR package uses the efficient fused LASSO algorithm which internally makes use of the Fused Lasso Signal Approximator (FLSA) algorithm [4]. The FLSA algorithm is an iterative algorithm that makes use of a Sub-Gradient Finding Algorithm (SFA). However, the original implementation of this algorithm in the MALSAR Matlab package is suitable only for linear time-series data. We extend this algorithm to handle general branching structure by re-implementing the Subgradient Finding Algorithm with gradient descent (**Algorithm 4**, $\text{SFA}_\text{G}$).

**Algorithm 1** is the main algorithm for the AGM method. The parameters $\alpha$ and $\gamma$, control the step size of AGM. The algorithm has two nested loops. The outer loop updates the step size parameters executing the main AGM framework. The inner loop uses the fused LASSO penalty to estimate new regression weights (**Algorithm 2**), which internally calls **Algorithm 3** (FLSA) and **Algorithm 4** ($SFA_G$) and updates the model parameters due to the non-smooth part.

**Algorithm 1:** DRMN-FUSED algorithm

**Input:**
- $\mathbf{Y}_{c,k}$ Gene expression vector for module $k$ in cell line $c$
- $\mathbf{X}_{c,k}$ Feature matrix for module $k$ in cell line $c$
- $\rho_1, \rho_2, \rho_3$
- $\Psi$ Lineage tree

**Output:**
- $W$ the resulting regression weights ($C \times F$, number of tasks by number of features)

$$W = \begin{pmatrix} W_1 \\ W_2 \\ \vdots \\ W_C \end{pmatrix}$$

**Initialze:**
Initialize regression weights for each task, $W_c = 0$, $W_c^{old} = 0$
$t = 1, t_{\text{old}} = 0, \gamma = 1, \gamma_{\text{inc}} = 2,$ ▷ Rate parameters for AGM
$F_{old} = 0$
**while** *not converged* **do**
 $\alpha = \frac{t_{\text{old}}-1}{t}$
 $W_c^{new} = (1+\alpha)W_c - \alpha W_c^{old}$
 $\nabla W_c^{new} = (\mathbf{X}_{c,k}W_c^{\intercal} - \mathbf{Y}_{c,k})^{\intercal}\mathbf{X}_{c,k}$ ▷ Compute gradient for the squared loss for all $c$
 $F_{new} = \sum_c \frac{1}{2}||\mathbf{Y}_{c,k} - \mathbf{X}_{c,k}W_c^{new\intercal}||_2^2$
 **while** *True* **do**
  $V_c = W_c^{new} - \nabla W_c^{new}/\gamma$ ▷ Get initial estimate of regression weight for all $c$
  $W^{proj}$ = getFGLASSO($V$, $\frac{\rho_1}{\gamma}$, $\frac{\rho_2}{\gamma}$, $\frac{\rho_3}{\gamma}$, $\Psi$) ▷ Fused Group LASSO projection (**Algorithm 2**)
  $F_{proj} = \sum_c \frac{1}{2}||\mathbf{Y}_{c,k} - \mathbf{X}_{c,k}W_c^{proj\intercal}||_2^2$
  $\Delta W^{proj} = W^{proj} - W^{new}$
  $F_\gamma = F_{new} + \sum_{i,j}(\Delta W^{proj} \odot \nabla W^{new})_{i,j} + \frac{\gamma}{2}||\Delta W^{proj}||_2^2$
  **if** $F_{proj} \leq F_\gamma$ **then**
   *break*
  **end**
  $\gamma = \gamma \times \gamma_{\text{inc}}$
 **end**
 $W_c^{old} = W_c$
 $W_c = W_c^{proj}$
 $F_{new} = \sum_c ||\mathbf{Y}_{c,k} - \mathbf{X}_{c,k}W_c^{\intercal}||_2^2 + \rho_1||W||_1 + \rho_2||\Psi W||_1 + \rho_3||W||_{2,1}$
 **if** $|F_{new} - F_{old}| <$ *tolerance* **then**
  converged=true
 **end**
 $F_{old} = F_{new}$
 $t_{old} = t$
 $t = \frac{1}{2}(1 + \sqrt{1+4t^2})$
**end**

**Algorithm 2:** getFGLASSO algorithm

**Input:**

- $V$: $C \times F$, matrix of initial regression weights, $C$ is the number of tasks, $F$ is the number of features)
- $\lambda_1, \lambda_2, \lambda_3$
- $\Psi$ Lineage tree

**Output:**

- $W^{proj}$ Fused Group LASSO projection

**for** $i = 1...F$ **do**

  $\mathbf{v} = V_{.,i}$ ▷ Column $i$ of matrix $V$

  $\mathbf{w} = \text{flsa}(\mathbf{v}, \lambda_1, \lambda_2, \Psi)$

  $\mathbf{w}_i = \frac{\max(||\mathbf{w}||_2 - \lambda_3, 0)}{||\mathbf{w}||_2} \times \mathbf{w}$

**end**

$W^{proj} = (\mathbf{w}_1, \mathbf{w}_2, ..., \mathbf{w}_F)$

**Algorithm 3:** flsa algorithm

**Input:**

- $\mathbf{v}$, $C \times 1$, column of estimate of regression weights, $C$ is the number of tasks
- $\lambda_1, \lambda_2$, regularization parameters of Fused Lasso
- $\Psi$ Lineage tree

**Output:**

- $\mathbf{w}$ updated estimate of regression weights based on Fused Lasso penalty

Solve $\Psi\Psi^\intercal \mathbf{z} = \Psi \mathbf{v}$ for $\mathbf{z}$

$z_{max} = ||\mathbf{z}||_\infty$ ▷ Get the max absolute value of $\mathbf{z}$

**if** $\lambda_2 \geq z_{max}$ **then**

  $t = \text{mean}(\mathbf{v})$

  $$t = \begin{cases} t - \lambda_1 & \text{if } t > \lambda_1 \\ t + \lambda_1 & \text{if } t < -\lambda_1 \\ 0 & otherwise \end{cases}$$ ▷ Soft-thresholding

  $\mathbf{w} = (t, t, ..., t)^\intercal$ ▷ $C \times 1$ vector

**end**

**else**

  $\mathbf{w} = \textbf{SFA}_G(\mathbf{v}, \lambda_2, \Psi, \mathbf{z})$

  $$\mathbf{w}(i) = \begin{cases} \mathbf{w}(i) - \lambda_1 & \text{if } \mathbf{w}(i) > \lambda_1 \\ \mathbf{w}(i) + \lambda_1 & \text{if } \mathbf{w}(i) < -\lambda_1 \\ 0 & otherwise \end{cases}$$ ▷ Soft-thresholding on all elements of $\mathbf{w}$

**end**

**Algorithm 4:** Subgradient Finding Algorithm with Gradient descent (SFA$_G$)

**Input:**

-$\mathbf{v}$, $\lambda_2$,$\mathbf{z}$

-$\Psi$: Lineage

**Output:**

- $\mathbf{w}$: updated estimate of regression weight

$L$=largest eigen value of $\Psi$

**while** *not converged* **do**

▷ Convergence determined by max iterations or the duality gap

$g = \Psi\Psi^{\mathsf{T}}\mathbf{z} - \Psi\mathbf{v}$ ▷ Compute gradient

$\mathbf{z} = \mathbf{z} - (g/L)$

$\mathbf{z}(i) = \begin{cases} \lambda_2 & \text{if } \mathbf{z}(i) > \lambda_2 \\ -1 * \lambda_2 & \text{if } \mathbf{z}(i) < -\lambda_2 \end{cases}$ ▷ Project $\mathbf{z}$ in the limit of $[-\lambda_2, \lambda_2]$

$\mathbf{s} = \Psi\Psi^{\mathsf{T}}\mathbf{z} - \Psi\mathbf{v}$ ▷ get the gradient again

$gap = \lambda_2||\mathbf{s}||_1 + < \mathbf{s}, \mathbf{z} >$ ▷ Get the duality gap

**if** *gap* $<$ *tolerance* **then**

convergence=true

**end**

**end**

$\mathbf{w} = \mathbf{v} - \Psi^{\mathsf{T}}\mathbf{z}$

#### 1.2 DRMN-ST

The structure prior version of DRMN is also implemented for each module across all cell types. At each iteration, the algorithm uses greedy hill climbing to find a regulator that when added, improves the overall score of the model. As mentioned in the main text, the prior term decomposes over inferred interactions $f \rightarrow k$ (meaning feature $f$ is in regulatory program of module $k$):

$$P(G_1, \cdots, G_C) = \prod_{f \rightarrow k} P(\mathbf{I}_{f \rightarrow k}) \tag{2}$$

where

$$P(\mathbf{I}_{f \rightarrow k}) = P(I^{root}_{f \rightarrow k}) \prod_{c' \rightarrow c \in \Psi} P(I^{c}_{f \rightarrow k} | I^{c'}_{f \rightarrow k}) \tag{3}$$

and $I^c_{f \rightarrow c}$ is 1 if $f \rightarrow c$ is selected for cell line $c$ and 0 otherwise.

The data likelihood can be written as

$$P(\mathbf{X}_{c,k} | \mathbf{R}_{c,k}, \mathbf{Y}_{c,k}) \tag{4}$$

where $\mathbf{X}_{c,k}$ is gene expression vector of module $k$ in cell line $c$, $\mathbf{Y}_{c,k}$ is the feature matrix of module $k$ in cell line $c$, and $\mathbf{R}_{c,k} =< G_{c,k}, \Theta_{c,k} >$ is the regulatory program of module $k$ consisting of $G_{c,k}$ the inferred interactions and $\Theta_{c,k}$ the corresponding regression coefficients. The data likelihood can be estimated as a conditional normal distribution

$$P(\mathbf{X}_{c,k} | \mathbf{R}_{c,k}, \mathbf{Y}_{c,k}) \sim \prod_{g \in \text{module} k} N \left( \sum_{f \in G_{c,k}} \theta_{c,k}(f) \mathbf{Y}_{c,k}(g, f), \Sigma_{x|G_{c,k}} \right) \tag{5}$$

and $\Sigma_{x|G_{c,k}}$ denotes conditional variance. **Algorithm 5** outlines the greedy hill climbing algorithm that is called in each iteration of EM algorithm (see Algorithm 1 in main text) for each module $k$. The parameter $maxIter$ is set to 5, meaning that at most 5 new regulators can be added to the regulatory program of a module in each iteration of the EM algorithm.

**Algorithm 5:** DRMN-ST algorithm

**Input:**

- $\mathbf{Y}_{c,k}$ Gene expression vector for module $k$ in cell line $c$
- $\mathbf{X}_{c,k}$ Feature matrix for module $k$ in cell line $c$
- $\Psi$ Lineage tree

**Output:**

- $\mathbf{R}_{c,k} =< G_{c,k}, \Theta_{c,k} >$ The updated regulatory program

```
for maxIter number of iterations do
    for each feature f do
        for each cell line c do
            scoreImprovement ← improvement of data likelihood when adding f to regulatory
              program of k
            if scoreImprovement > 0 then
                I^c_{f→k} ← 1
            end
            else
                I^c_{f→k} ← 0
            end
        end
        Calculate P(G_1, ··· , G_C)                ▷ Prior term, see equations 2 and 3
        Calculate P(X_{c,k}|R_{c,k}, Y_{c,k})      ▷ Data likelihood term, see equations 4 and 5
    end
    Add f*, feature with highest overall score improvement to the model.
end
```

### 2 Supplementary Figures

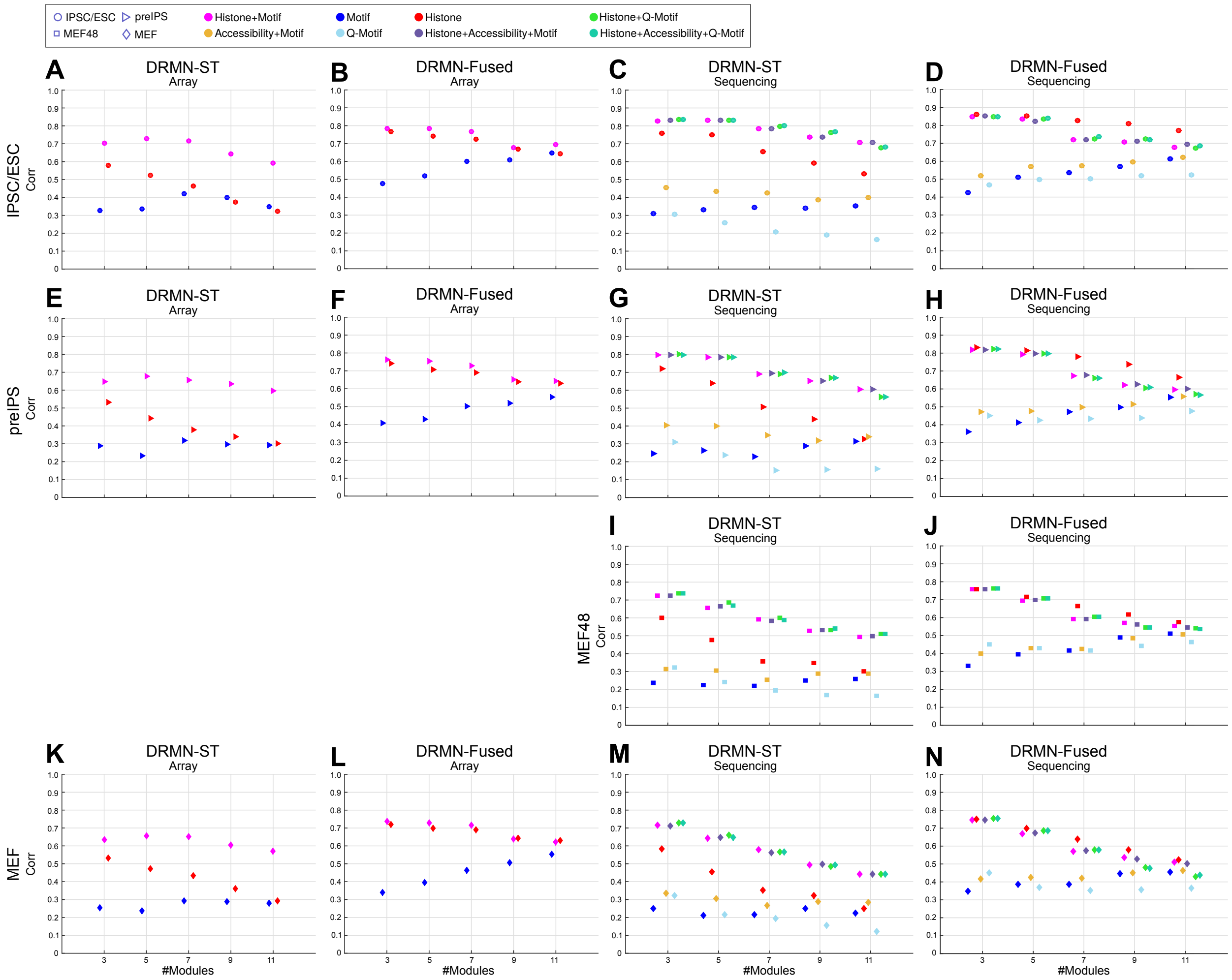

IPSC/ESC
preIPS
MEF48
MEF
Histone+Motif
Accessibility+Motif
Motif
Q-Motif
Histone
Histone+Accessibility+Motif
Histone+Q-Motif
Histone+Accessibility+Q-Motif
A DRMN-ST Array
B DRMN-Fused Array
C DRMN-ST Sequencing
D DRMN-Fused Sequencing
E DRMN-ST Array
F DRMN-Fused Array
G DRMN-ST Sequencing
H DRMN-Fused Sequencing
I DRMN-ST Sequencing
J DRMN-Fused Sequencing
K DRMN-ST Array
L DRMN-Fused Array
M DRMN-ST Sequencing
N DRMN-Fused Sequencing
Corr
#Modules

**Supplemental Figure S1.** Average per-module correlation as a function of different number of modules for DRMN and RMN models. Each row and marker shape corresponds to a cell line from the array or sequencing dataset. Different colors correspond to different feature types. Results in the main paper **Fig. 2** is only for the ESC/iPSC cell line. **A-D.** ESC/IPSC. **E-H.** pre-IPSC. **I, J.** MEF 48 hours (only for sequencing dataset). **K-N** MEF. Each column corresponds to DRMN-ST or DRMN-FUSED versions on a dataset (**A**, **E**, **K** for DRMN-ST on array dataset, **B**, **F**, **L** for DRMN-FUSED on array dataset, **C**, **G**, **I**, **M** for DRMN-ST on sequencing dataset, and **D**, **H**, **J**, **N** for DRMN-FUSED on sequencing dataset).

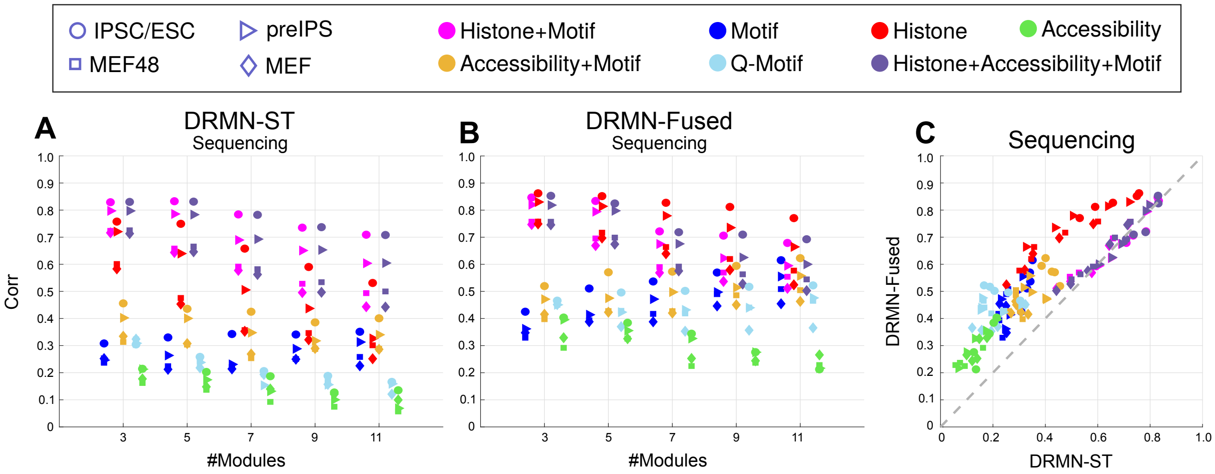
IPSC/ESC
preIPS
MEF48
MEF
Histone+Motif
Accessibility+Motif
Motif
Q-Motif
Histone
Accessibility
Histone+Accessibility+Motif
A
DRMN-ST
Sequencing
Corr
#Modules
B
DRMN-Fused
Sequencing
#Modules
C
Sequencing
DRMN-Fused
DRMN-ST

**Supplemental Figure S2.** Performance of Accessibility (ATAC-seq) as a single feature (green) vs. Motif (blue), Histone+Motif (magenta), Q-Motif (light blue), ATAC+Motif (yellow), Histone (red), and Histone+Accesibility+Motif (dark purple) using DRMN-ST (**A**) and DRMN-FUSED (**B**)). **C**. Scatter plot comparing DRMN-ST vs. DRMN-Fused for the same features.

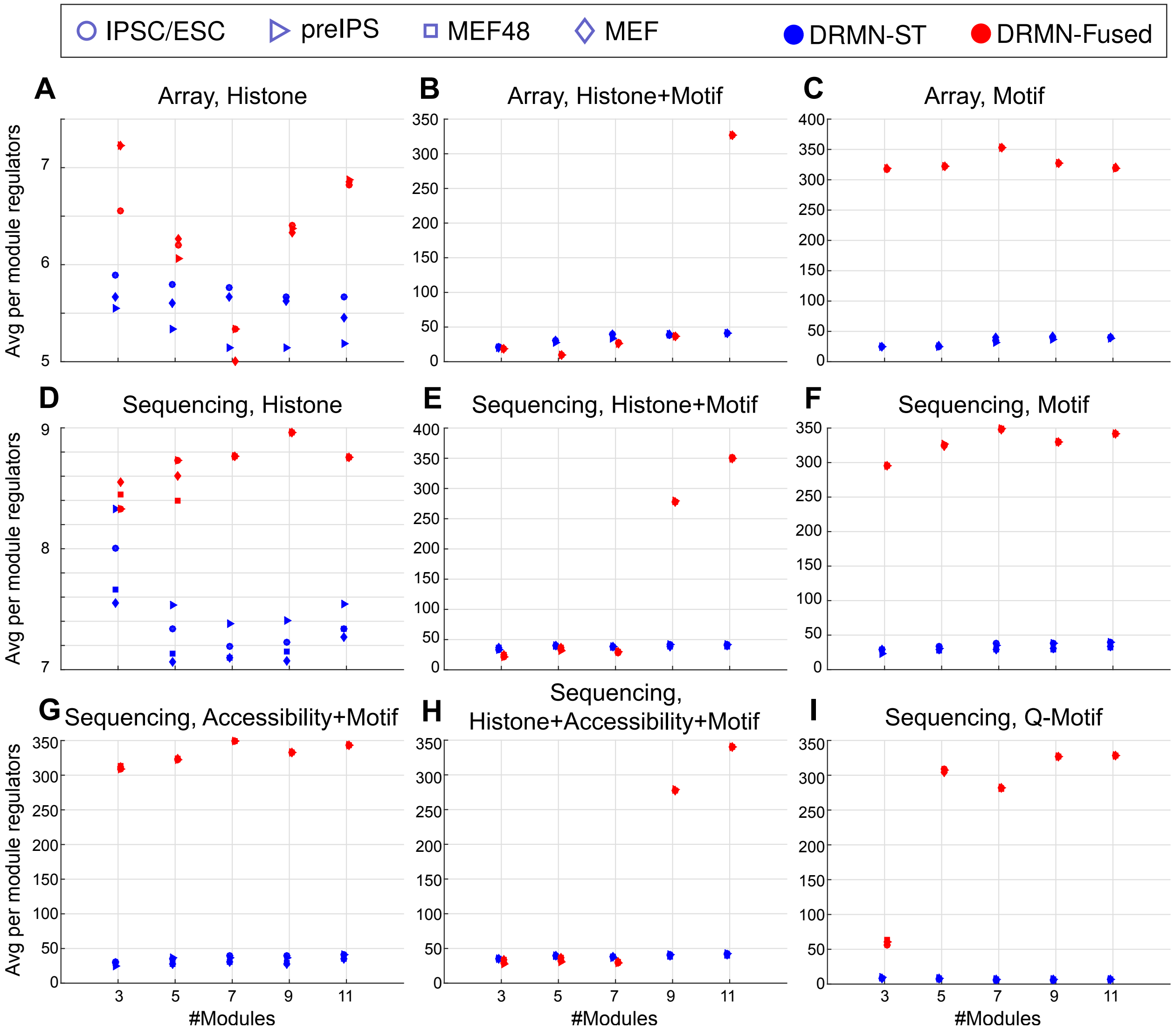
IPSC/ESC
preIPS
MEF48
MEF
DRMN-ST
DRMN-Fused
A
Array, Histone
B
Array, Histone+Motif
C
Array, Motif
D
Sequencing, Histone
E
Sequencing, Histone+Motif
F
Sequencing, Motif
G
Sequencing, Accessibility+Motif
H
Sequencing, Histone+Accessibility+Motif
I
Sequencing, Q-Motif
Avg per module regulators
#Modules

**Supplemental Figure S3.** Average number of regulators per module of DRMN-ST and DRMN-FUSED models for different dataset/feature combinations **A-C.** Feature sets in the array dataset. **D-I.** Feature sets in the sequencing dataset.

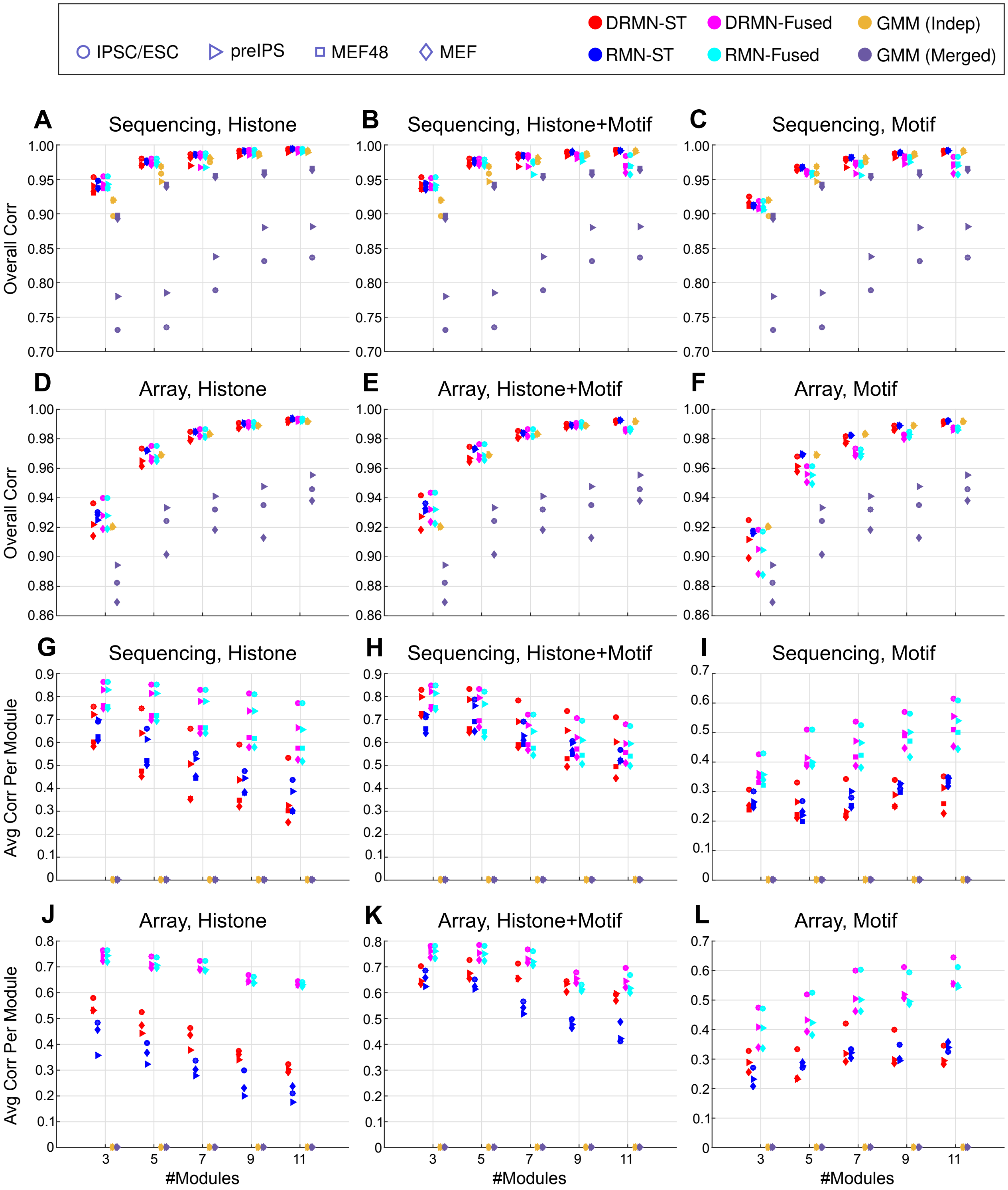

IPSC/ESC
preIPS
MEF48
MEF
DRMN-ST
DRMN-Fused
GMM (Indep)
RMN-ST
RMN-Fused
GMM (Merged)
A Sequencing, Histone
B Sequencing, Histone+Motif
C Sequencing, Motif
D Array, Histone
E Array, Histone+Motif
F Array, Motif
G Sequencing, Histone
H Sequencing, Histone+Motif
I Sequencing, Motif
J Array, Histone
K Array, Histone+Motif
L Array, Motif
Overall Corr
Avg Corr Per Module
#Modules

**Supplemental Figure S4.** Comparing DRMN's expression predictions to baseline approaches. Each panel compares six algorithms on the basis of one correlation metric (y-axis) across a range of $k$ (x-axis) for a particular feature type. Each dot represents the results for one cell type averaged over three-fold cross validation. **A-F**. Comparison based on Pearson correlation of predicted to true expression for held-aside genes. **G-L**. Comparison based on per-module correlation coefficients, averaged across $k$ modules.

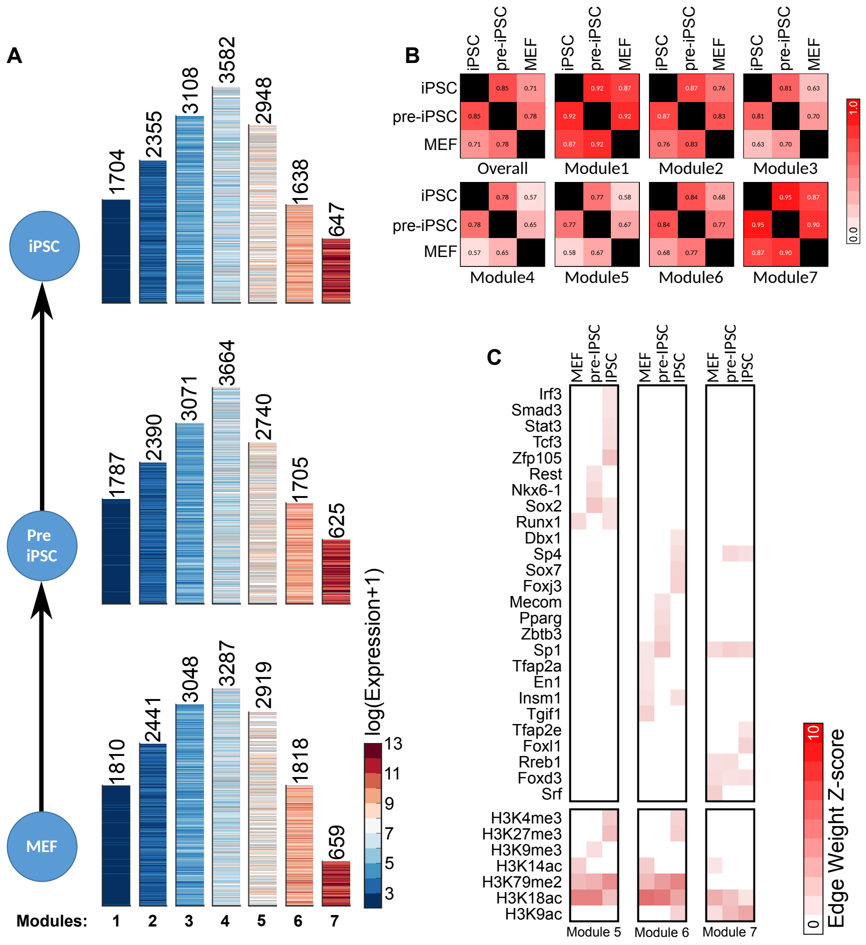
A
iPSC
Pre iPSC
MEF
1704
2355
3108
3582
2948
1638
647
1787
2390
3071
3664
2740
1705
625
1810
2441
3048
3287
2919
1818
659
Modules: 1 2 3 4 5 6 7
log(Expression+1)
13
11
9
7
5
3
B
iPSC
pre-iPSC
MEF
Overall
Module1
Module2
Module3
Module4
Module5
Module6
Module7
1.0
0.0
C
MEF
pre-iPSC
iPSC
Irf3
Smad3
Stat3
Tcf3
Zfp105
Rest
Nkx6-1
Sox2
Runx1
Dbx1
Sp4
Sox7
Foxj3
Mecom
Pparg
Zbtb3
Sp1
Tfap2a
En1
Insm1
Tgif1
Tfap2e
Foxl1
Rreb1
Foxd3
Srf
H3K4me3
H3K27me3
H3K9me3
H3K14ac
H3K79me2
H3K18ac
H3K9ac
Module 5
Module 6
Module 7
Edge Weight Z-score
10
0

**Supplemental Figure S5.** Application of DRMNs to the cellular reprogramming array dataset using histone marks and motifs for the feature set. **A**. Show are gene expression patterns of the $k = 7$ modules. The number above the heatmap is the number of genes in that module. **B**. Similarity of modules across cellular stages as measured by F-score. The color intensity is proportional to the match. **C**. Inferred regulators for the three most highly expressed modules across cell stages.

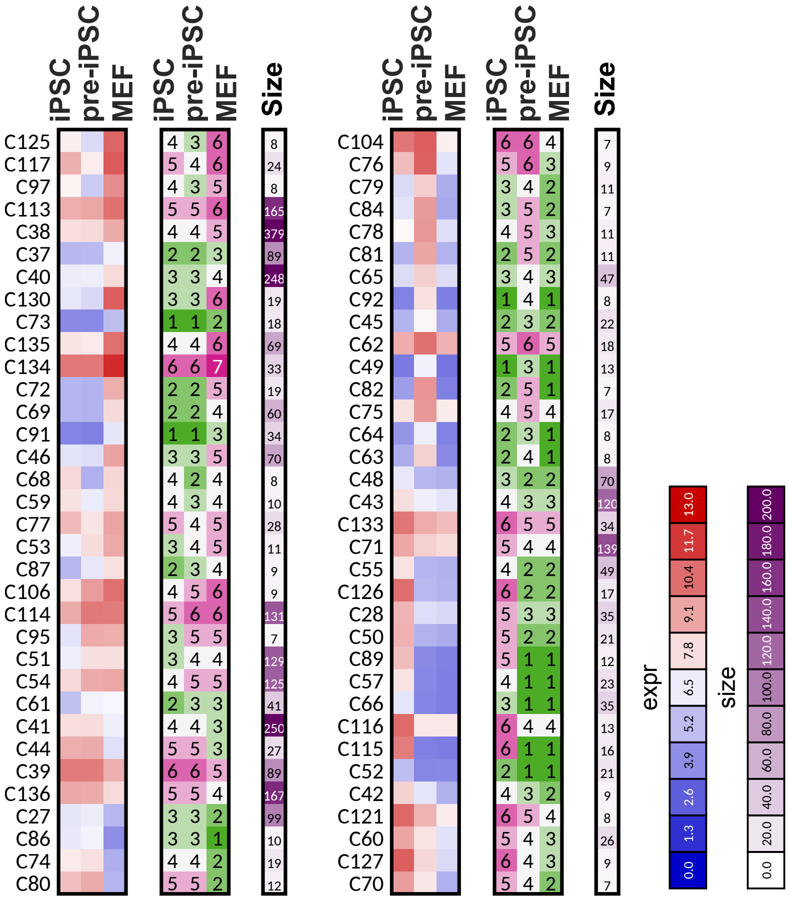
iPSC
pre-iPSC
MEF
iPSC
pre-iPSC
MEF
Size
C125 C117 C97 C113 C38 C37 C40 C130 C73 C135 C134 C72 C69 C91 C46 C68 C59 C77 C53 C87 C106 C114 C95 C51 C54 C61 C41 C44 C39 C136 C27 C86 C74 C80
C104 C76 C79 C84 C78 C81 C65 C92 C45 C62 C49 C82 C75 C64 C63 C48 C43 C133 C71 C55 C126 C28 C50 C89 C57 C66 C116 C115 C52 C42 C121 C60 C127 C70
expr
0.0 1.3 2.6 3.9 5.2 6.5 7.8 9.1 10.4 11.7 13.0
size
0.0 20.0 40.0 60.0 80.0 100.0 120.0 140.0 160.0 180.0 200.0

**Supplemental Figure S6.** Transitioning genes sets for the array dataset. Shown on left is the average expression of the genes in the group, in the middle are module assignments, and on the right are the number of genes in that group.

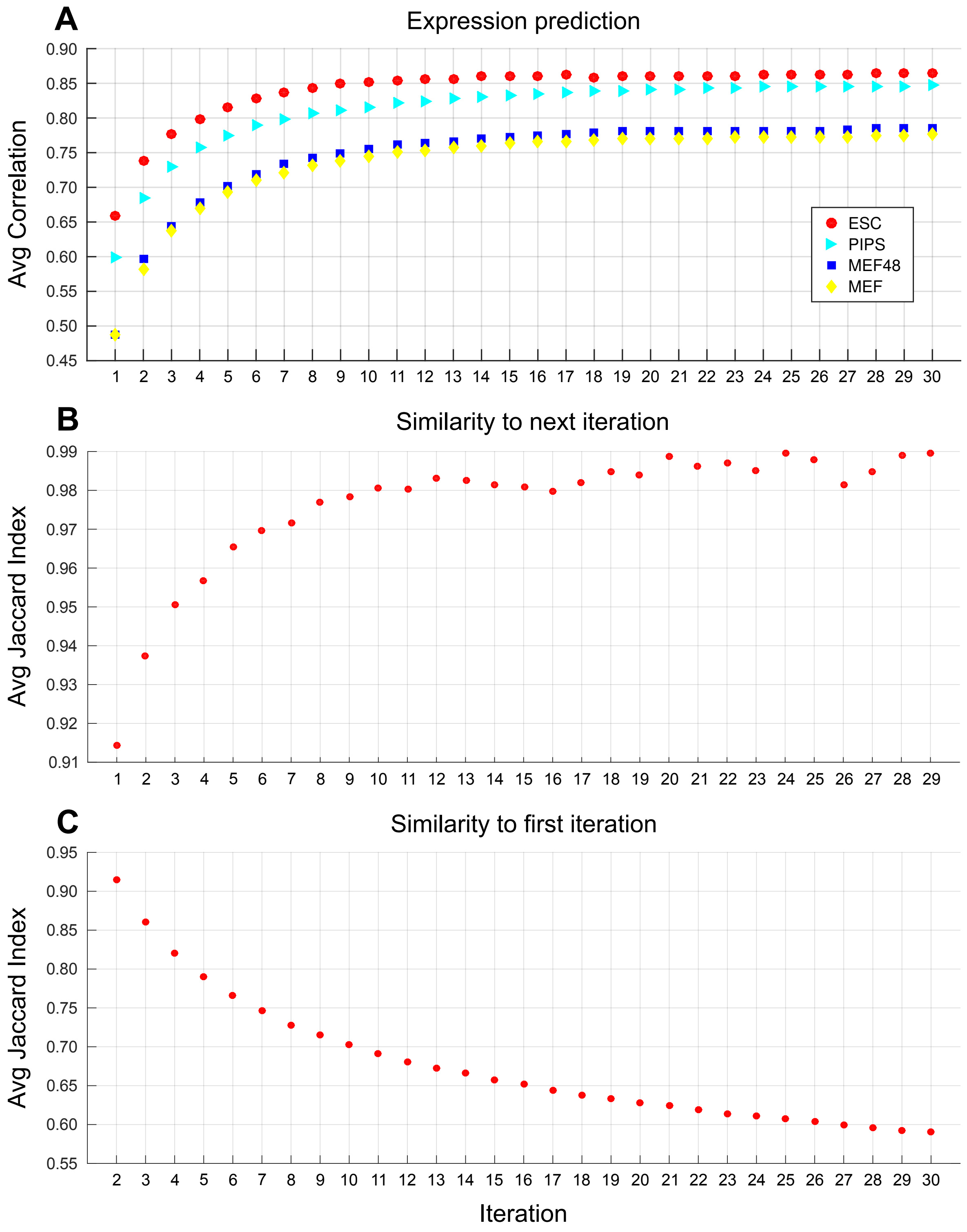
A
Expression prediction
Avg Correlation
ESC
PIPS
MEF48
MEF
B
Similarity to next iteration
Avg Jaccard Index
C
Similarity to first iteration
Avg Jaccard Index
Iteration

**Supplemental Figure S7.** The effect of number of iterations on DRMN performance. The results are with Motif+Chromatin, k=3, using fused LASSO on the reprogramming sequencing dataset ($\rho_1 = 100, \rho_2 = 50, \rho_3 = 0$). **A**. Average correlation (over all modules) for each cell line, as a function of number of iterations. Each marker corresponds to a cell line. **B**. Similarity of module assignments between consecutive iterations used to assess the stability and convergence of results. **C**. Similarity of module assignments between iteration 1 and iteration $i$.

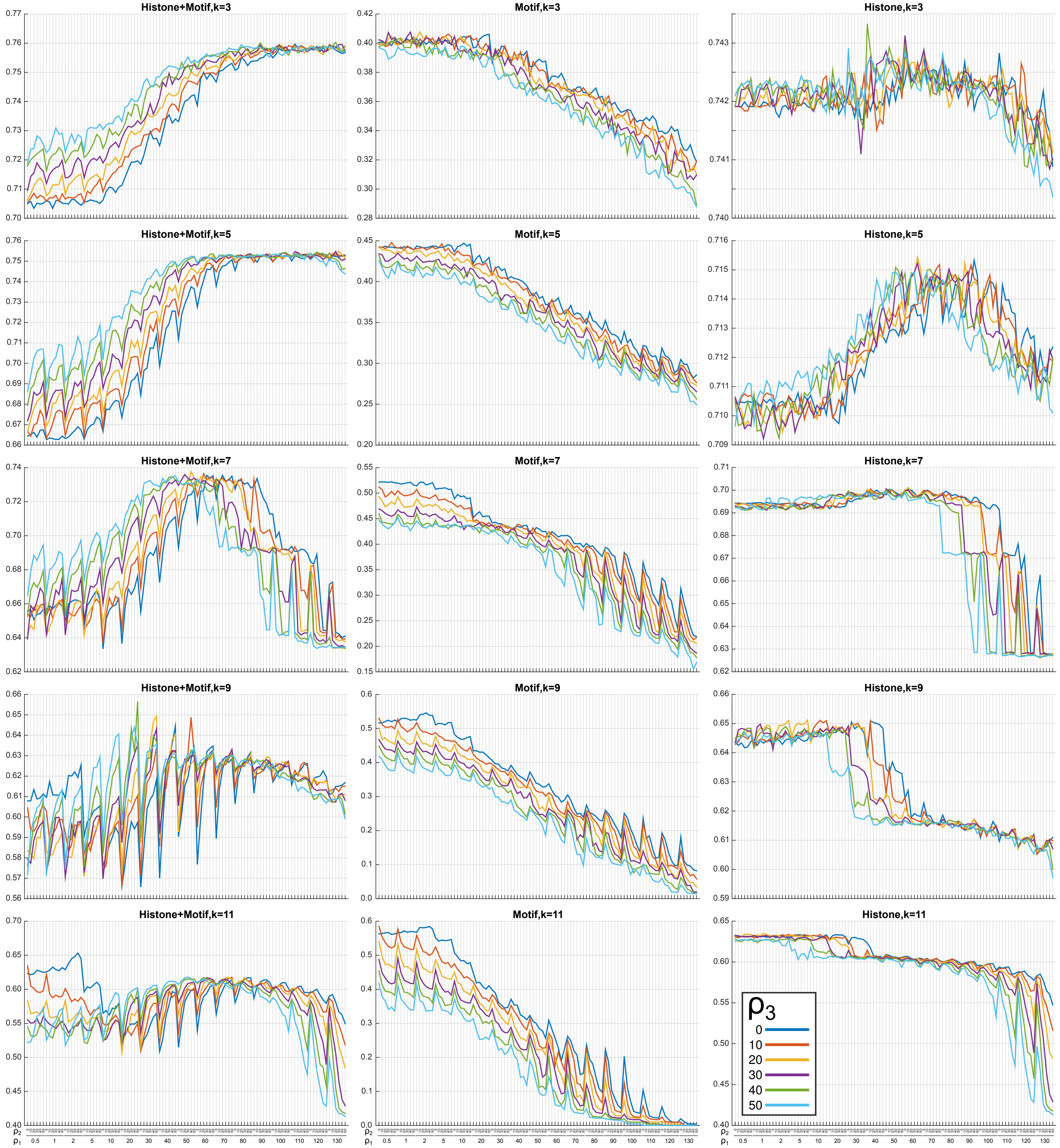
Histone+Motif,k=3
Motif,k=3
Histone,k=3
Histone+Motif,k=5
Motif,k=5
Histone,k=5
Histone+Motif,k=7
Motif,k=7
Histone,k=7
Histone+Motif,k=9
Motif,k=9
Histone,k=9
Histone+Motif,k=11
Motif,k=11
Histone,k=11
$\rho_3$
0
10
20
30
40
50
$\rho_2$
$\rho_1$
0.5 1 2 5 10 20 30 40 50 60 70 80 90 100 110 120 130

**Supplemental Figure S8.** The effect of hyper-parameters on DRMN performance for Histone+Motif, Histone, and Motif in array dataset. Each column corresponds to type of feature sets and rows correspond to module sizes. In each panel, different colors corresponds to different values of $\rho_3$, the group LASSO penalty (enforcing selection of same features for all cell lines).

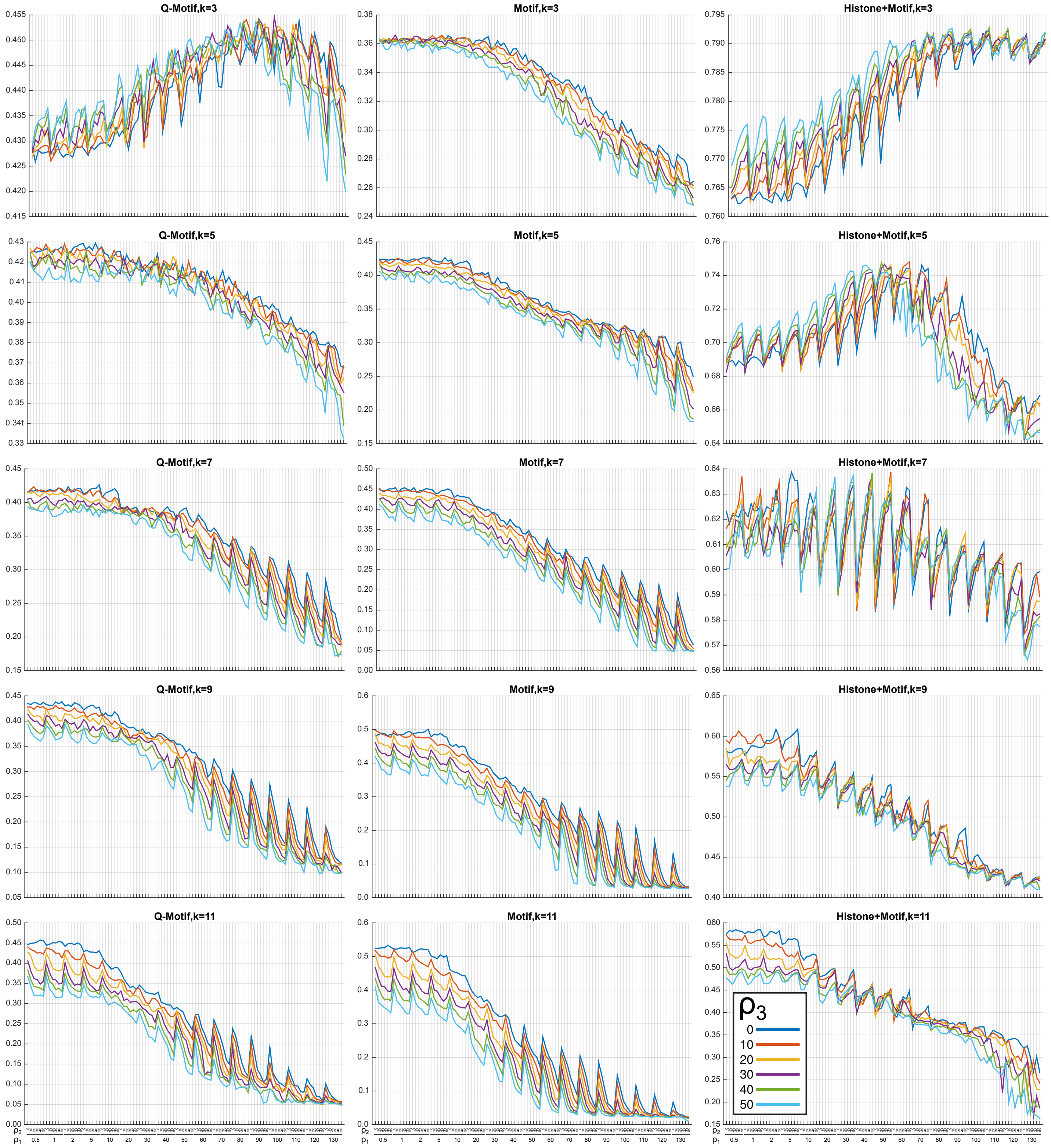
Q-Motif,k=3
Motif,k=3
Histone+Motif,k=3
Q-Motif,k=5
Motif,k=5
Histone+Motif,k=5
Q-Motif,k=7
Motif,k=7
Histone+Motif,k=7
Q-Motif,k=9
Motif,k=9
Histone+Motif,k=9
Q-Motif,k=11
Motif,k=11
Histone+Motif,k=11
$\rho_3$
0
10
20
30
40
50
$\rho_2$
$\rho_1$
0.5 1 2 5 10 20 30 40 50 60 70 80 90 100 110 120 130

**Supplemental Figure S9.** The effect of hyper-parameters on DRMN performance for Q-Motif, Motif, and Histone+Motif features. Each column corresponds to type of feature sets and rows correspond to module sizes. In each panel, different colors corresponds to different values of $\rho_3$, the group LASSO penalty (enforcing selection of same features for all cell lines).

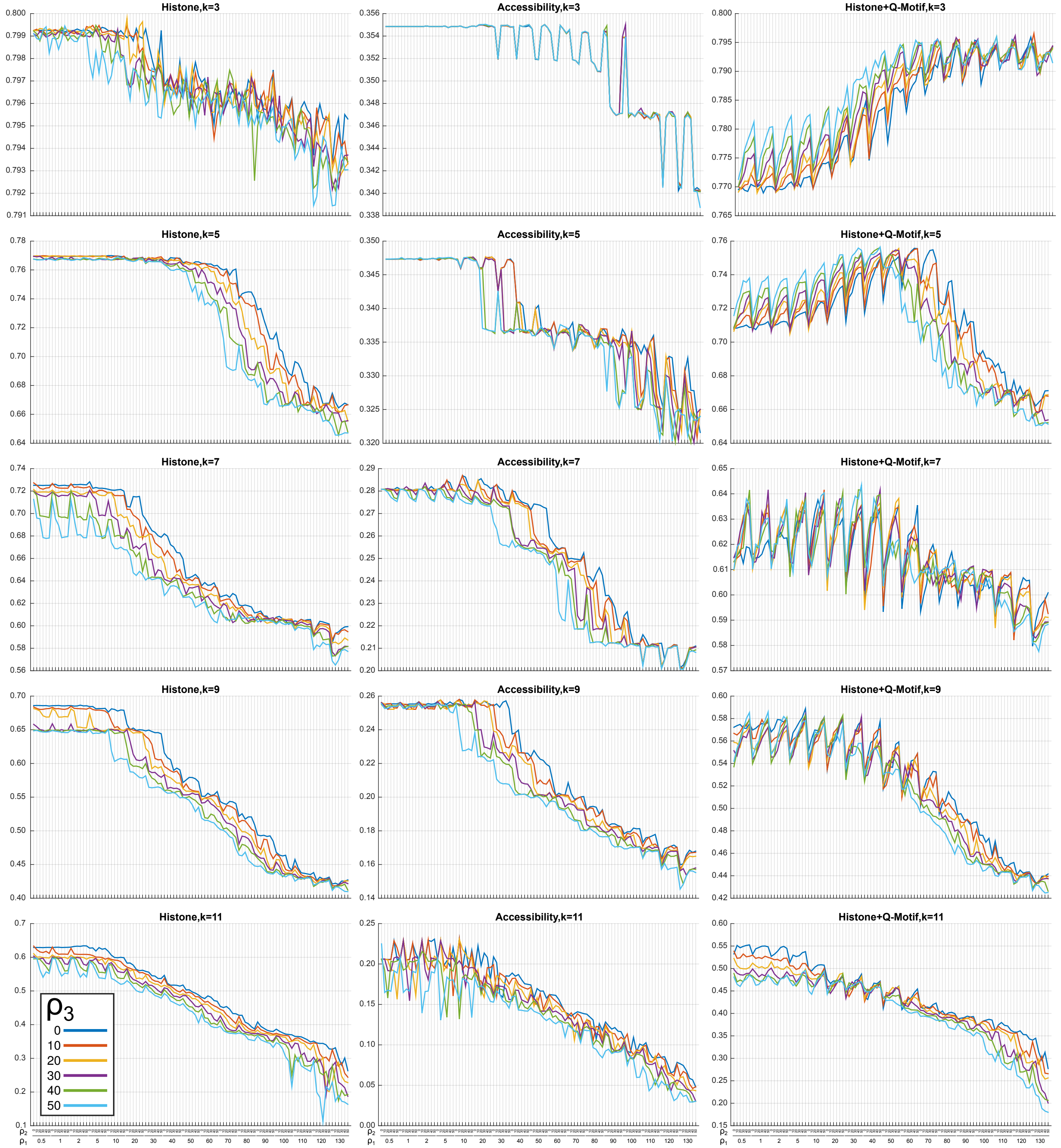
Histone,k=3
Accessibility,k=3
Histone+Q-Motif,k=3
Histone,k=5
Accessibility,k=5
Histone+Q-Motif,k=5
Histone,k=7
Accessibility,k=7
Histone+Q-Motif,k=7
Histone,k=9
Accessibility,k=9
Histone+Q-Motif,k=9
Histone,k=11
Accessibility,k=11
Histone+Q-Motif,k=11
ρ3
0
10
20
30
40
50
ρ2
ρ1
0.5 1 2 5 10 20 30 40 50 60 70 80 90 100 110 120 130

**Supplemental Figure S10.** The effect of hyper-parameters on DRMN performance for Histone, Accessibility, and Histone+Q-Motif features. Each column corresponds to type of feature sets and rows correspond to module sizes. In each panel, different colors corresponds to different values of $\rho_3$, the group LASSO penalty (enforcing selection of same features for all cell lines).

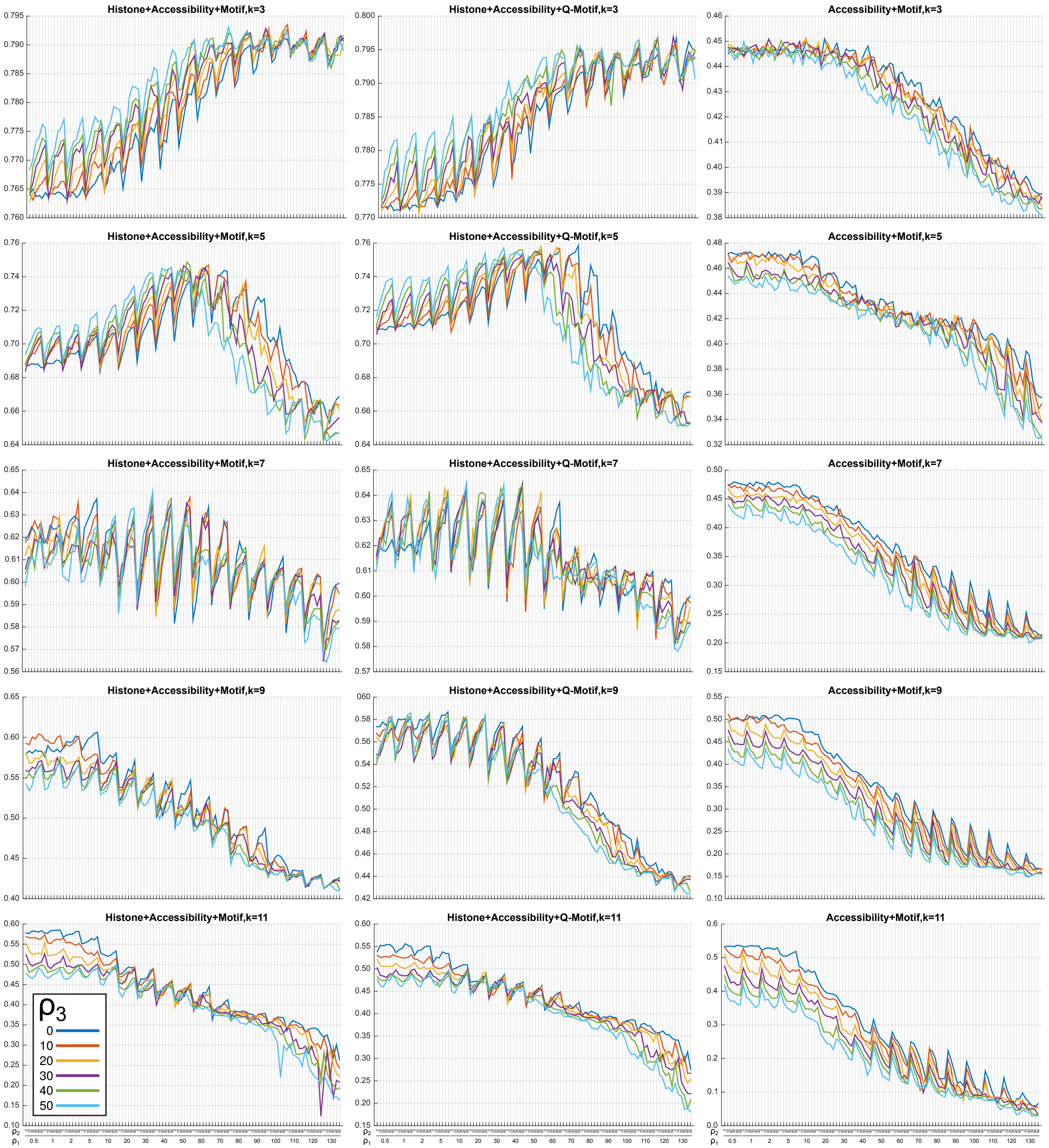
Histone+Accessibility+Motif,k=3
Histone+Accessibility+Q-Motif,k=3
Accessibility+Motif,k=3
Histone+Accessibility+Motif,k=5
Histone+Accessibility+Q-Motif,k=5
Accessibility+Motif,k=5
Histone+Accessibility+Motif,k=7
Histone+Accessibility+Q-Motif,k=7
Accessibility+Motif,k=7
Histone+Accessibility+Motif,k=9
Histone+Accessibility+Q-Motif,k=9
Accessibility+Motif,k=9
Histone+Accessibility+Motif,k=11
Histone+Accessibility+Q-Motif,k=11
Accessibility+Motif,k=11
ρ3
0
10
20
30
40
50
ρ2
ρ1
0.5 1 2 5 10 20 30 40 50 60 70 80 90 100 110 120 130

**Supplemental Figure S11.** The effect of hyper-parameters on DRMN performance for Histone+Accessibility+Motif, Histone+Accessibility+Q-Motif, and Accessibility+Motif. Each column corresponds to type of feature sets and rows correspond to module sizes. In each panel, different colors corresponds to different values of $\rho_3$, the group LASSO penalty (enforcing selection of same features for all cell lines).

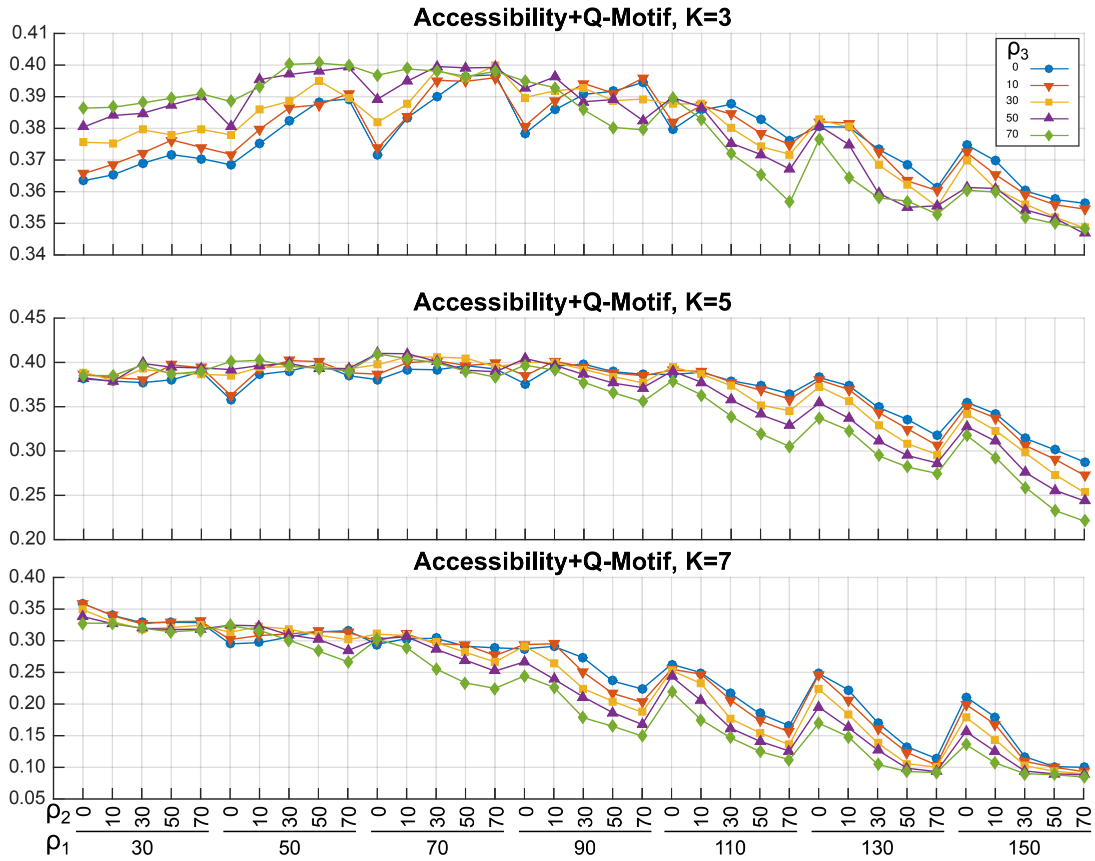

Accessibility+Q-Motif, K=3
Accessibility+Q-Motif, K=5
Accessibility+Q-Motif, K=7
ρ3
0
10
30
50
70
ρ2
ρ1
30
50
70
90
110
130
150

**Supplemental Figure S12.** Hyper-parameter selection for the dedifferentiation dataset. Rows correspond to different number of modules. In each panel, different colors corresponds to different values of $\rho_3$, the group LASSO penalty (enforcing selection of same features for all cell lines).

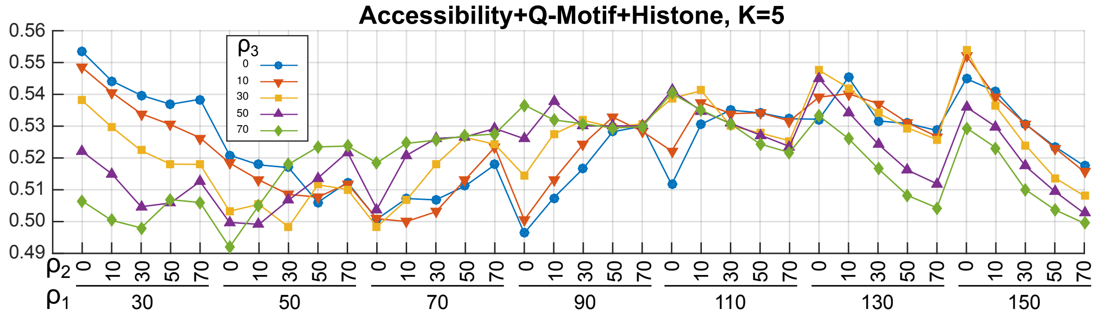
Accessibility+Q-Motif+Histone, K=5
$\rho_3$
0
10
30
50
70
0.56
0.55
0.54
0.53
0.52
0.51
0.50
0.49
$\rho_2$
$\rho_1$
30
50
70
90
110
130
150

**Supplemental Figure S13.** Hyper-parameter selection for the ESC differentiation dataset. Different colors correspond to different values of $\rho_3$, the group LASSO penalty (enforcing selection of same features for all cell lines).

### References

[1] Jiayu Zhou, Jianhui Chen, and Jieping Ye. Malsar: Multi-task learning via structural regularization – users manual version 1.1, 2012.

[2] Yu. Nesterov. Smooth minimization of non-smooth functions. *Mathematical Programming*, 103(1):127–152, May 2005.

[3] Yu. Nesterov. Gradient methods for minimizing composite objective function. CORE Discussion Papers 2007076, Universit catholique de Louvain, Center for Operations Research and Econometrics (CORE), 2007.

[4] Jun Liu, Lei Yuan, and Jieping Ye. An efficient algorithm for a class of fused lasso problems. In *Proceedings of the 16th ACM SIGKDD International Conference on Knowledge Discovery and Data Mining*, KDD '10, page 323332, New York, NY, USA, 2010. Association for Computing Machinery.
